# Diverse *Fgfr1* signaling pathways and endocytic trafficking regulate early mesoderm development

**DOI:** 10.1101/2024.02.16.580629

**Authors:** James F. Clark, Philippe Soriano

## Abstract

The Fibroblast growth factor (FGF) pathway is a conserved signaling pathway required for embryonic development. Activated FGF receptor 1 (FGFR1) drives multiple intracellular signaling cascade pathways, including ERK/MAPK and PI3K/AKT, collectively termed canonical signaling. However, unlike *Fgfr1* null embryos, embryos containing hypomorphic mutations in *Fgfr1* lacking the ability to activate canonical downstream signals are still able to develop to birth, but exhibit severe defects in all mesodermal-derived tissues. The introduction of an additional signaling mutation further reduces the activity of *Fgfr1,* leading to earlier lethality, reduced somitogenesis, and more severe changes in transcriptional outputs. Genes involved in migration, ECM-interaction, and phosphoinositol signaling were significantly downregulated, proteomic analysis identified changes in interactions with endocytic pathway components, and cells expressing mutant receptors show changes in endocytic trafficking. Together, we identify processes regulating early mesoderm development by mechanisms involving both canonical and non-canonical *Fgfr1* pathways, including direct interaction with cell adhesion components and endocytic regulation.

## Introduction

Fibroblast Growth Factor (FGF) signaling plays an integral role in development, driving numerous cellular processes including proliferation, differentiation, and cellular adhesion (Clark and Soriano, 2022; Ornitz and Itoh, 2022). FGFs are a family of secreted proteins that bind to and activate their cognate FGF receptors (FGFRs), which are receptor tyrosine kinases (RTKs). Binding of an FGF to its receptor results in the dimerization of FGFRs and subsequent transphosphorylation of multiple tyrosine residues of the FGFRs. Upon activation, FGFRs recruit multiple effectors to these modified residues to engage downstream intracellular signaling pathways, including ERK/MAPK and PI3K/AKT (Fig. 1A) (Brewer et al., 2016). The mammalian FGF signaling family consists of fifteen canonical FGF ligands and four canonical FGFRs (Ornitz and Itoh, 2015, 2022).

**Figure 1.**
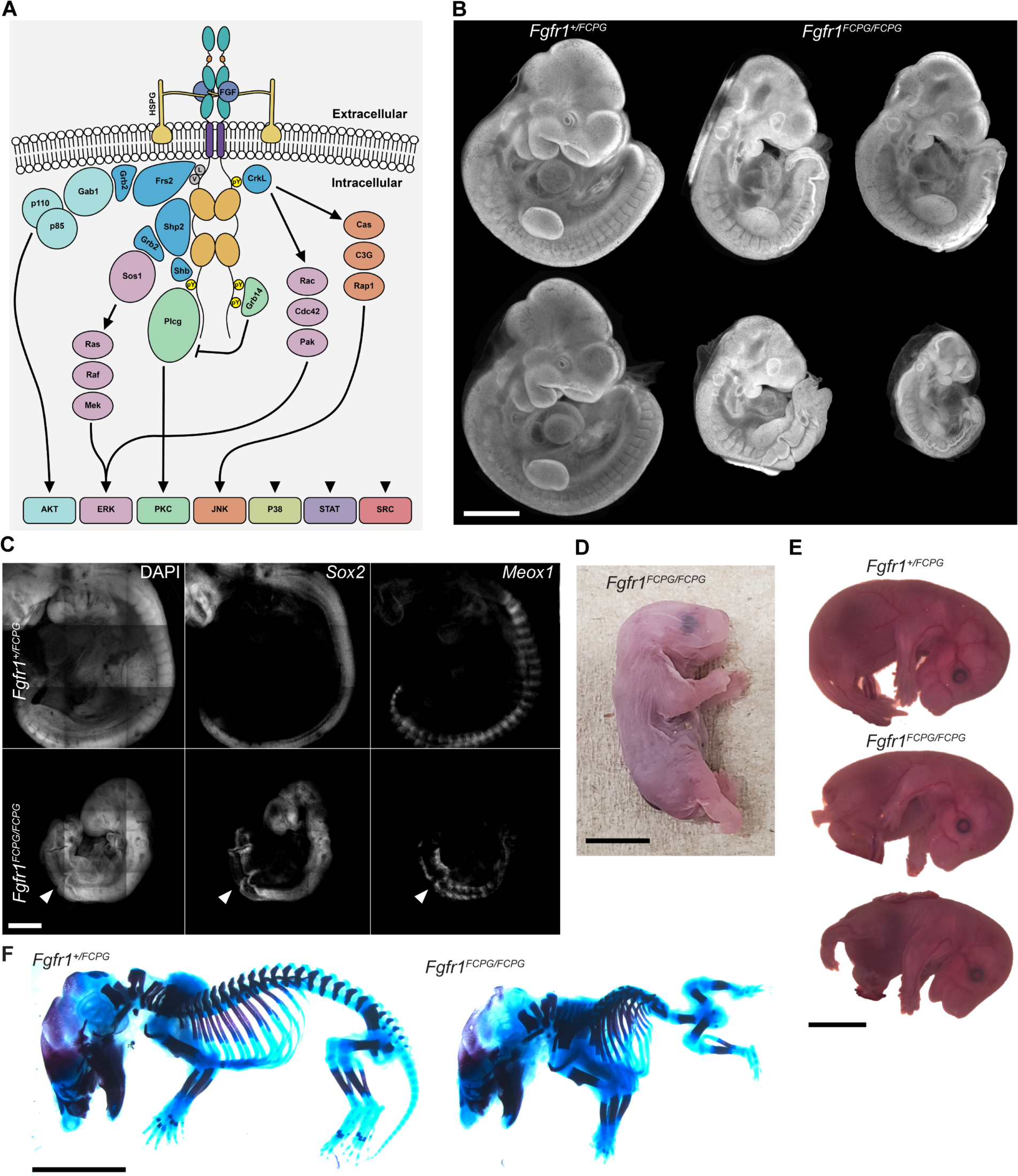
A portion of homozygous *Fgfr1^FCPG^* mutant mice survive perinatally while exhibiting severe axial truncation. **A)** Activated FGFRs engage multiple downstream intracellular signaling pathways through the recruitment of various effector proteins to its intracellular domain. **B)** DAPI staining of E10.5 embryos. *Fgfr1^FCPG/FCPG^*mutants display reduced size and craniofacial, somite, and neural tube defects, as previously reported. However, variance in the severity of the phenotypes ranges between individuals. Scale bar represents 1mm. **C)** At E10.5, *Fgfr1^FCPG/FCPG^* embryos display disrupted somite patterning and early truncation based on *Meox1* expression. The neural tube is formed, as highlighted with *Sox2* expression, but does not close in all mutant embryos. Scale bar represents 50μm. **D)** *Fgfr1^FCPG/FCPG^* P0 pups display several phenotypes including axial truncation, *spina bifida*, hindlimb malformation, and lack a tail. Scale bar represents 5mm. **E)** *Fgfr1^FCPG/FCPG^* embryos collected at E14.5 exhibit spina bifida, hindlimb malformation, body wall closure defects, and axial truncation. Scale bar represents 2mm. **F)** Skeletal staining of E14.5 wild type and homozygous *Fgfr1^FCPG^*embryos. Mutant embryos exhibit multiple severe skeletal defects including vertebral truncation, rib fusions, abnormal digit patterning, pelvic bone agenesis, and reduced ossification. Scale bar represents 2mm.

Both *Fgfr1* and *Fgfr2* are necessary for early mouse development (Deng et al., 1994; Xu et al., 1998; Yamaguchi et al., 1994; Yu et al., 2003). Deletion of *Fgfr1*, on a 129S4 genetic background, results in embryonic lethality at peri-implantation while deletion of *Fgfr2* results in lethality at mid-gestation (Brewer et al., 2015; Kurowski et al., 2019; Molotkov et al., 2017). Modulation of the FGF signaling pathway during embryonic development has identified multiple roles in various stages of somitic, intermediate, and lateral plate mesoderm. In the growing embryo, the neuromesodermal progenitors (NMPs) in the tailbud serve as the progenitor pool for the posterior extension of the neural tube and mesoderm. FGF, Notch, and Wnt signals overlap and integrate to maintain pluripotency in the tailbud (Wahl et al., 2007). As the body axis elongates, the balance of signals is shifted, resulting in subsequent specification of the neural and mesodermal lineages (Déqueant et al., 2006; Déqueant and Pourquié, 2008; Mongera et al., 2019).

Chemical perturbation of FGFRs in the chick tailbud results in arrested expansion of the somitic mesoderm (Delfini et al., 2005). Additionally, previous data has shown that deletion of some FGF ligands results in vertebral phenotypes. Loss of *Fgf4* or *Fgf8* in the presomitic mesoderm resulted in multiple defects in the axial skeleton, with loss of *Fgf4* presenting a greater degree of disruption (Anderson et al., 2020). *Fgf3* null mice also exhibit truncated tails with reduced caudal vertebrae (Anderson et al., 2016). In addition, previous analysis of hypomorphic *Fgfr1* alleles identified homeotic transformations in thoracic vertebrae (Brewer et al., 2015; Partanen et al., 1998).

Alongside the somitic mesoderm, FGF signaling plays a role in intermediate mesoderm development, giving rise to both renal and gonadal tissues (Davidson et al., 2019). Both *Fgfr1* and *Fgfr2* have known roles in renal morphogenesis. Conditional deletion of either *Fgfr1* or *Fgfr2* in the metanephric mesenchyme, using *Pax3Cre*, yielded normal kidneys, however combined deletion of both receptors resulted in rudimentary ureteric bud formation at E11.5 (Poladia et al., 2006). *Hoxb7Cre* deletion of *Fgfr1* in the ureteric bud resulted in no phenotype, while deletion of *Fgfr2* resulted in disrupted branching (Zhao et al., 2004). Similarly, embryos that lack *Fgf9* and *Fgf20* form ureteric buds that begin to grow into the metanephric mesenchyme but fail to branch, and embryos that lack *Fgf8* exhibit deficiencies in nephron development (Barak et al., 2012; Perantoni et al., 2005).

FGF signaling also contributes to the development of the lateral plate mesoderm which flanks the intermediate mesoderm and gives rise to parts of the viscera, body wall, and limb buds (Prummel et al., 2020). During limb bud growth, reciprocal FGF signaling between the limb mesenchyme and apical ectodermal ridge (AER) regulates outgrowth (Ornitz and Marie, 2019). Both *Fgf8* and *Fgf10* null mice exhibit severe limb truncations (Lewandoski et al., 2000; Moon and Capecchi, 2000; Sekine et al., 1999). *Fgfr2* null mice display complete lack of limb bud formation at E10.5 (Molotkov et al., 2017; Xu et al., 1998). Conversely, conditional *Fgfr1* null mice develop limb buds but exhibit reduced limb growth and digit patterning defects (Li et al., 2005; Verheyden et al., 2005).

We have been dissecting how these FGFRs engage signaling to drive developmental processes by introducing point mutations that ablate the recruitment of specific effectors to the receptor. This allows us to isolate the role of individual or groups of downstream signals important for developmental processes. The most severe combinatorial *Fgfr1* and *Fgfr2* alleles to date, *Fgfr^FCPG^*(that eliminates binding of <u>F</u>RS2, <u>C</u>RKL/II, SHB/<u>P</u>LCγ, <u>G</u>RB14), resulted in the loss of all FGF-induced ERK1/2, AKT, PLCγ, STAT, and a significant reduction of p38 and JNK activation (Brewer et al., 2015; Ray et al., 2020). We collectively refer to these signal transduction cascades as canonical FGF signaling given their well-established functions downstream of FGFRs. Embryos expressing the FCPG alleles of either *Fgfr1* or *Fgfr2* survived well beyond the peri-implantation lethality seen in *Fgfr1* null animals (Brewer et al., 2015; Kurowski et al., 2019) or the mid-gestation lethality seen in *Fgfr2* null animals (Molotkov et al., 2017).

This raises the question of what functionality remains in the FCPG alleles and how these functions are engaged outside of canonical FGF signaling. Our previous results indicated that while FGFR1/2 regulate cell-matrix and cell-cell adhesion, these processes were not affected in *Fgfr1^FCPG^* or *Fgfr2^FCPG^* homozygous embryos (Ray et al., 2020; Ray and Soriano, 2023). Evidence suggests that both FGFR1 and FGFR2 can recruit SRC family kinases (SFKs). SFKs are known to function downstream of multiple receptor tyrosine kinases, are involved in the promotion of cell proliferation and differentiation, and are critical for integrin signaling (Klinghoffer et al., 1999; Li et al., 2004; Sandilands et al., 2007). Previous evidence suggests that FGFR1 can directly recruit SFKs through the phosphorylation of Y730 (Dudka et al., 2010). FGFR2 also interacts with SFKs, although a direct binding site has not been identified (Schuller et al., 2008). However, phosphorylation of the equivalent site in FGFR2 (Y734) was instead shown to recruit PIK3R2, the p85 regulatory subunit of PI3K (Salazar et al., 2009). FGFR1 and FGFR2 are known to activate PI3K, however the recruitment of PIK3R2 to FGFR2 is involved in endocytic recycling of the receptor independent of PI3K/AKT activation (Francavilla et al., 2013). These results suggest that Y730/Y734 may be a multifunctional site linking FGFR1/2 to SFKs and endocytic trafficking. Together, these data indicate that various effectors may still be binding to FGFR1/2^FCPG^ to provide functionality beyond canonical signaling, which we refer to as non-canonical FGF activity.

To investigate which developmental processes depend on canonical FGFR1 signaling, we first examined the developmental defects present in *Fgfr1^FCPG/FCPG^*embryos. One third of these embryos survive perinatally, albeit with severe developmental defects. Using RNA-seq analysis, we find significant changes in genes involved in ECM interactions and Wnt signaling. *Fgfr1^FCPG/FCPG^*embryos also exhibit defects in all mesodermal lineages, with disruption of intermediate mesoderm formation including complete kidney agenesis, during late gastrulation and early embryogenesis. To address the question of residual function, we next produced a novel *Fgfr1* allele, *Fgfr1^FCSPGMyc^*, containing an additional mutation to eliminate proposed interactions with SFKs and endocytic recycling. Homozygous *Fgfr1^FCSPGMyc^* embryos display more severe phenotypes than *Fgfr1^FCPGMyc^* mutants but still do not phenocopy the null allele. Using additional RNA-seq analysis we identify two key pathways, ECM-receptor interactions and phosphatidylinositol signaling, that may account for the phenotypic differences seen between the two signaling mutant embryos. Proteomic analysis and proximity ligation assays further reveal that FGFR1 is able to complex with FAK and SFKs, indicating possible recruitment to focal adhesions. Additionally, multiple proteins involved in endocytic recycling were identified as potential FGFR1 interactors, including PI3KR2. Colocalization of FGFR1 to either early or late endosomes highlighted differences in endocytic trafficking between FGFR1^WTMyc^, FGFR1^FCPGMyc^, and FGFR1^FCSPGMyc^. Together, these data indicate that *Fgfr1* maintains considerable functionality when lacking canonical signaling, at least in part through recruitment to focal adhesions and endocytic trafficking.

## Results

### Fgfr1^FCPG/FCPG^ mutants survive past E10.5 and display severe mesodermal defects

As previously published, the *Fgfr1^FCPG^* allele prevents all canonical downstream signal activation through multiple pathways, including MAPK/ERK and PI3K/AKT (Brewer et al., 2015; Ray et al., 2020). To investigate what developmental functions can proceed in the absence of canonical signaling, and to facilitate these studies, we generated an *Fgfr1^FCPGMyc^*allele that contained the previous *FCPG* mutations along with a 2x Myc tag at the C-terminus. Similarly, we generated an *Fgfr1^Myc^* allele to determine the effects of the 2x Myc tag on wild-type *Fgfr1* function. *Fgfr1^Myc/Myc^*mice are viable and fertile, with no apparent phenotypes (data not shown), demonstrating that the peptide tags appended to the C-terminus do not compromise the activity of the receptors. Subsequent analyses focus on *Fgfr1^FCPGMyc^*, referred to hereafter as *Fgfr1^FCPG^* for simplicity.

As previously observed, homozygous *Fgfr1^FCPG^* embryos exhibited multiple defects including reduced size, loss of the second pharyngeal arch, improper somite patterning, and a kinked neural tube (Fig. 1B). At E10.5, *Fgfr1^FCPG/FCPG^* embryos exhibited disruption to both neural tube closure and somite patterning, as marked by *Sox2* and *Meox1*, respectively, via *in situ* Hybridization Chain Reaction (HCR). Proper morphology was maintained until about the level of the first limb bud. Posterior to the first limb bud, somites were hypomorphic and unevenly spaced, and the overlying neural tube was buckled, a phenotype often associated with mesodermal extension defects (Fig. 1C). We did not observe the presence of any ectopic neural tube tissue nor excess neural epithelium through whole-mount *in situ* or histological sections.

Analysis of the original *Fgfr1^FCPG^* strain (Brewer et al., 2015) indicated that homozygous embryos were not recoverable past E11.5. However, during our investigations we recovered some dead *Fgfr1^FCPG/FCPG^* mutants at birth, and verified this result across three independent lines, the original *Fgfr1^FCPG^* line and two *Fgfr1^FCPGMyc^* lines, to rule out any background effects of a single line. Upon further investigation, we found that one third of homozygous *Fgfr1^FCPG/FCPG^* embryos survive past gastrulation (Table 1).

Embryos that continue developing past E10.5 have beating hearts with normal vascularization of the amniotic sac. These data suggest that a threshold level of canonical FGF signaling is required for development at or around E10.5; sufficient signaling allows the embryo to continue developing, while insufficient signaling results in arrested development and necrosis of the embryo. P0 homozygous mutant pups recovered at birth exhibited severe axial truncations, with complete lack of a tail, hind limb malformations, digit patterning defects on both fore- and hind-limbs, closed *spina bifida*, and dorsal blood clots (Fig. 1D). Phenotypes consistent with those seen at E10.5 were observed in E14.5 embryos and skeletal staining revealed reduced ossification and multiple skeletal abnormalities (Fig. 1E, F). Multiple vertebrae along the posterior were malformed or missing, digit patterning defects were observed in the fore- and hind-limbs, the pelvic bones were hypomorphic, and the cranial vault was underdeveloped.

To determine which tissues require *Fgfr1* signaling, we utilized published single-cell RNAseq data to look for tissues with high expression of *Fgfr1* before and after E10.5 (Cao et al., 2019; Pijuan-Sala et al., 2019). We utilized these datasets to identify cell populations with high *Fgfr* expression and at what timepoints the *Fgfr1* signaling mutations might have the greatest effect. We found that *Fgfr1* showed the highest expression from E9.5 to E11.5. However, *Fgfr2* was more highly expressed than *Fgfr1* at E9.5 and E10.5, while *Fgfr3* and *Fgfr4* were expressed in very few cells at all stages (Fig. 2A). The expression of *Fgfr1* from E9.5 to E11.5 aligned with the phenotypes we see in our *Fgfr1^FCPG^* embryos, indicating that there might be an integral requirement for *Fgfr1* function at this time, with expression highest in mesodermal tissues (Fig. 2A, Fig. S1A, B).

**Figure 2.**
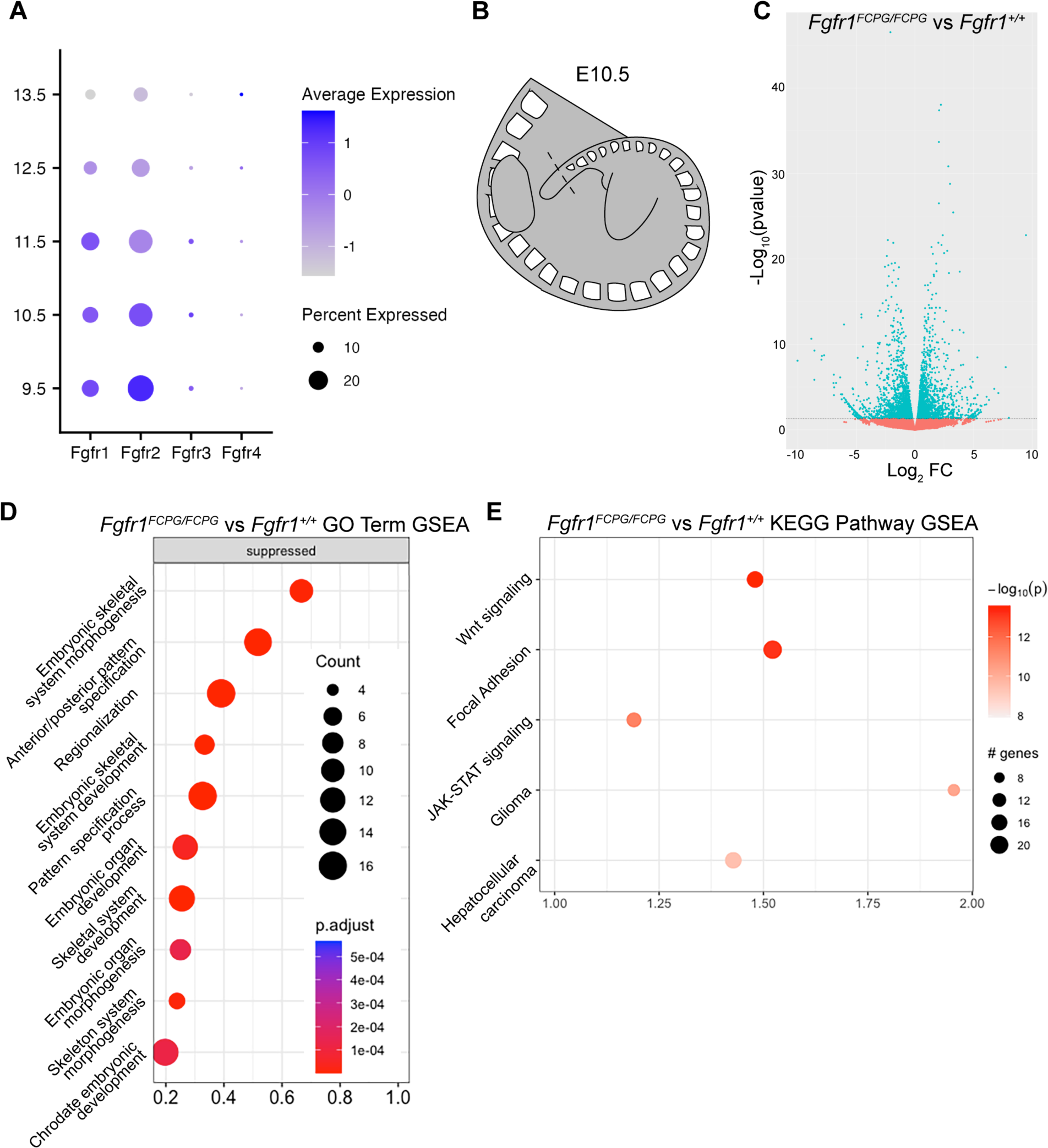
RNA-seq analysis of *Fgfr1^FCPG^* tailbud tissue reveals downregulation of genes involved mesodermal patterning, Wnt signaling, and cell adhesion. **A)** Analysis of a whole-embryo scRNA-seq organogenesis dataset (Cao et al., 2019; Pijuan-Sala et al., 2019) reveals high expression of both *Fgfr1* and *Fgfr2* from E9.5 to E11.5. **B)** Bulk RNA-seq analysis was performed using tailbud tissue dissected from E9.5, E10.5, and E11.5 embryos. Tailbud tissue was cut at the posterior-most definitive somite boundary (dashed lines). **C)** Volcano plots displaying differential gene expression between *Fgfr1^FCPG/FCPG^*mutants and wild type at E10.5. Horizontal dashed line signifies significance cut-off of p=0.05. **D)** GO Term GSEA of down-regulated genes in *Fgfr1FCPG/FCPG* tailbuds at E10.5. The most significant terms involve skeletal morphogenesis, anterior-posterior patterning, and regionalization. **E)** KEGG Pathway GSEA of down-regulated genes in *Fgfr1^FCPG/FCPG^* tailbuds at E10.5. Mutants exhibit changes in ECM-receptor interaction and focal adhesion genes as well as multiple signaling pathways including Wnt and JAK-STAT.

The high expression of *Fgfr2* at E9.5 and E10.5 suggests that it could compensate for impaired *Fgfr1^FCPG^* signaling by the formation of heterodimers (Bellot et al., 1991; Ueno et al., 1992). Because FGFR heterodimers have never been observed endogenously, we created a line carrying a 3 x FLAG tag at the C terminus of the wild-type *Fgfr2* (*Fgfr2^Flag^*) to use in conjunction with the *Fgfr1^Myc^* allele (Fig. S2A). *Fgfr2^Flag/Flag^*, and *Fgfr1^Myc/Myc^; Fgfr2^Flag/Flag^* mice are viable and fertile, with no apparent phenotypes (data not shown), demonstrating that as for FGFR1, the peptide tag appended to the C-terminus does not compromise the activity of FGFR2. We then used the *Fgfr1^Myc^* and *Fgfr2^Flag^*alleles in a proximity ligation assay (PLA) to indeed detect the presence of FGFR1:FGFR2 heterodimers in somitic mesoderm (Fig. S2B). However, while heterodimer formation might be a factor underlying the severity of the *Fgfr1^FCPG/FCPG^* phenotype, previous data indicates that heterodimers are not responsible for the continued development of the FCPG alleles compared to the null, as *Fgfr1^FCPG/FCPG^;Fgfr2^FCPG/FCPG^* embryos survived to E8.5 and exhibit normal expression of mesodermal markers, while *Fgfr1^-/-^;Fgfr2^-/-^* embryos die at implantation (Kurowski et al., 2019; Ray et al., 2020). The presence of heterodimers in wild-type tissue reinforces this point, indicating that the discrepancy between the FCPG and null alleles is more likely due to inherent function in the hypomorphic allele, beyond canonical signaling, rather than through dimerization with other FGFRs.

### Fgfr1^FCPG/FCPG^ mutants have changes in somitic and intermediate mesoderm gene expression

As FGF signaling has known roles in NMP maintenance and somitogenesis we next examined developmental progression of the tailbud (Delfini et al., 2005; Déqueant et al., 2006; Déqueant and Pourquié, 2008; Mongera et al., 2019; Wahl et al., 2007). To determine what transcriptional changes were being induced by the loss of canonical FGF signaling in the NMP and somitic mesoderm, we performed bulk RNA-seq on tailbuds of wild-type and *Fgfr1^FCPG/FCPG^* embryos at E10.5. Tailbud tissue was excised just caudal to the youngest definitive somite to capture the pool of NMPs alongside differentiating populations proximal to the tailbud (Fig. 2B). RNA-seq identified a number significantly differentially expressed genes (DEGs) (Fig. 2D, Table 2). Gene ontology (GO) Term Biological Process (BP) Gene Set Enrichment Analysis (GSEA) identified several processes involved in patterning of the embryo, mainly pertaining to anterior-posterior patterning and skeletal system development (Fig. 2D).

Many of the most significantly downregulated genes were Hox family members (Wellik, 2007), as well as *Meox1*, a key regulator of somitic mesoderm (Mankoo et al., 2003), and *Pax2*, an important regulator of intermediate mesoderm development (Torres et al., 1995) (Table 2). As expression of these regulators of patterning and specification was reduced, it appears that the *Fgfr1^FCPG^* mutation functions upstream to regulate these processes, consistent with both previously published data and the phenotypes observed in the mutant embryos (Déqueant and Pourquié, 2008; Mongera et al., 2019). Kyoto Encyclopedia of Genes and Genomes (KEGG) Pathway GSEA identified Focal Adhesion, Wnt, and JAK-STAT signaling pathways as the most significantly altered in *Fgfr1^FCPG/FCPG^* mutants (Fig. 2E, Table 3). These data indicate that loss of canonical FGF activity alters regulation of Wnt signaling, cell proliferation, and cell adhesion, all of which have documented interactions with *Fgfrs* (Goto et al., 2017; Ohkubo et al., 2004; Ray et al., 2020; Ray and Soriano, 2023; Wahl et al., 2007).

### Fgfr1^FCPG/FCPG^ embryos exhibit defects in vertebrae and digit patterning

To determine how these expression changes manifest in the observed phenotypes, we examined later stage *Fgfr1^FCPG^*embryos. Initial observation of homozygous *Fgfr1^FCPG^* mutant physical abnormalities revealed truncated axis, hind limb malformations, and digit patterning at P0. RNA-seq analysis revealed significant disruption to genes involved in skeletal development (Fig. 3A). Previous studies found that perturbation of FGF activity in somitic tissues resulted in vertebral defects (Anderson et al., 2020; Anderson et al., 2016; Brewer et al., 2015; Partanen et al., 1998). Accordingly, we observed multiple skeletal defects in *Fgfr1^FCPG/FCPG^* mutants. Axial truncation in the skeleton was detected from E14.5 onward, with multiple missing vertebrae in the lumbar and sacral regions. Malformation of the pelvic bones was also observed, with some mutants showing almost complete agenesis of the pelvis, which altered the formation of the hind limbs (Fig. 3B, Fig. S3A). As the pelvic bones are part of the appendicular skeleton derived from the lateral plate mesoderm, disrupted pelvic morphology indicates that loss of canonical FGF activity also impacts lateral plate mesoderm development in addition to somitic and intermediate mesoderm.

**Figure 3.**
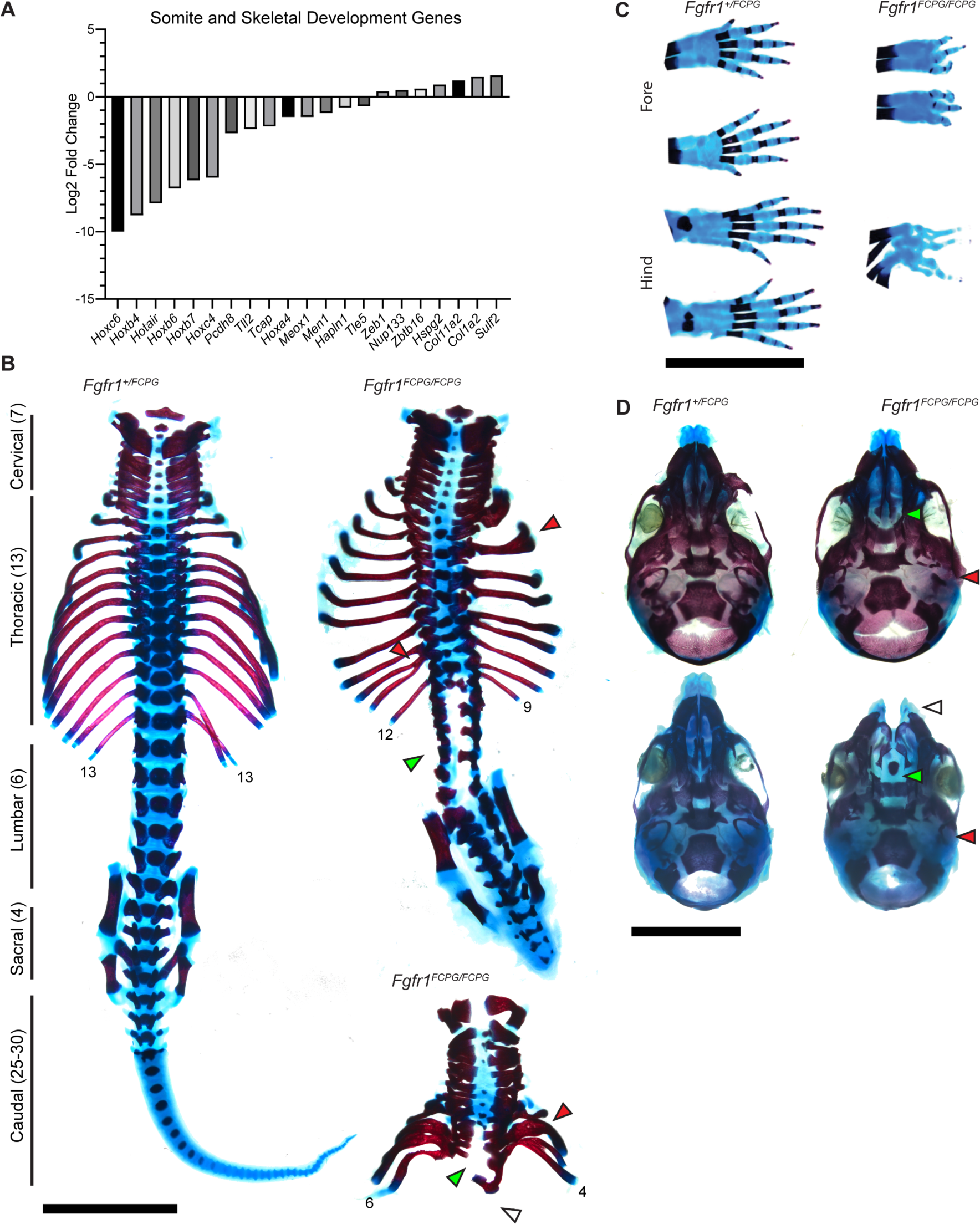
*Fgfr1^FCPG^*allele reveals extensive skeletal morphology and ossification defects up to perinatal development. **A)** RNA-seq of *Fgfr1^FCPG^* embryos indicates a large downregulation of genes involved in somite and skeletal development compared to wild type. Key genes, such as multiple Hox genes and Meox1, are downregulated while certain extracellular matrix genes are slightly upregulated. **B)** A wild-type P0 spinal column with vertebral groupings labeled. Homozygous *Fgfr1^FCPG^* embryos exhibit a range of axial truncation severity, as depicted by two P0 spinal columns (right). Red arrows denote examples of rib fusions, green arrows depict regions of *spina bifida*, and white arrow depicts complete lack of sacral vertebrae and pelvic bones. Scale bar represents 5mm. **C)** Digit patterning defects are seen in both the fore and hind paws, as well as delayed ossification of the digits. Fusion of paws was observed rarely (n = 1/14). Scale bar represents 5mm. **D)** Craniofacial skeletal defects include reduced ossification throughout the cranial vault, loss of the occipital bone (red arrow), and cleft palate (green arrow) (n = 13/14) or face (white arrow) (n = 1/14). Scale bar represents 5mm.

Skeletal staining also revealed multiple fusions of the digits in both the fore and hind limbs (Fig. 3C, Fig. S3B). The number of digits formed in the *Fgfr1^FCPG/FCPG^* mutants ranged from three to four on each limb, with disrupted patterning of the carpals and metacarpals. Interestingly, this indicates a major function for non-canonical *Fgfr1* signaling in digit patterning and limb formation. Loss of FGF signaling has known defects in proximal distal patterning of the developing limb (Mariani et al., 2008). However, our data indicate non-canonical signaling is sufficient for patterning and formation of the limb (Fig. 1F). Conversely, canonical signaling is required for proper patterning of the digits. Previous studies have focused predominantly on the roles of Shh and BMP signaling in digit patterning, with FGF signaling acting downstream of these pathways during digit tip formation (Sanz-Ezquerro and Tickle, 2003; Tickle, 2006). Additionally, the digits showed a severe lack of or delay in ossification. Ossification delay was observed as well in the skull, with multiple bones showing reduced bone deposition (Fig. 3D). *Fgfr1^FCPG^* mutants also consistently lacked proper formation of the occipital bone and exhibit cleft palate, with some embryos (1/13) having a more severe cleft face. These results are more severe than previously published data using an *Fgfr1^FCPG^* allele over a conditional *Fgfr1* knock-out in the neural crest (Brewer et al., 2015). Together, these data reveal that loss of canonical signaling downstream of *Fgfr1* does not recapitulate many of the skeletal defects associated with conditional knockout of *Fgfr1*, highlighting a greater role for non-canonical signaling.

### Fgfr1^FCPG/FCPG^ embryos exhibit intermediate mesoderm defects and kidney agenesis

Because we observed disruption to genes relating to intermediate mesoderm development via RNA-seq (Fig. 4A, Table 2), we next wanted to determine how the intermediate mesoderm-derived tissues were affected in *Fgfr1^FCPG^* mutants. Strikingly, we observed complete and bilateral kidney agenesis in E18.5 *Fgfr1^FCPG/FCPG^* embryos during dissections for skeletal preparations. As both *Fgfr1* and *Fgfr2* have known roles in renal morphogenesis we opted to examine ureteric bud formation (Barak et al., 2012; Poladia et al., 2006; Sims-Lucas et al., 2011; Zhao et al., 2004). Renal morphogenesis begins at E8.5 with the formation of the intermediate mesoderm and the appearance of the nephric duct and the nephrogenic chord. By E9.5, epithelialization of the nephric duct occurs and the metanephric mesenchyme starts to condense. At E10.5, the nephric duct buds into the metanephric mesenchyme with full ureteric bud formation by E11.0, which begins branching at E11.5 (Costantini and Kopan, 2010).

**Figure 4.**
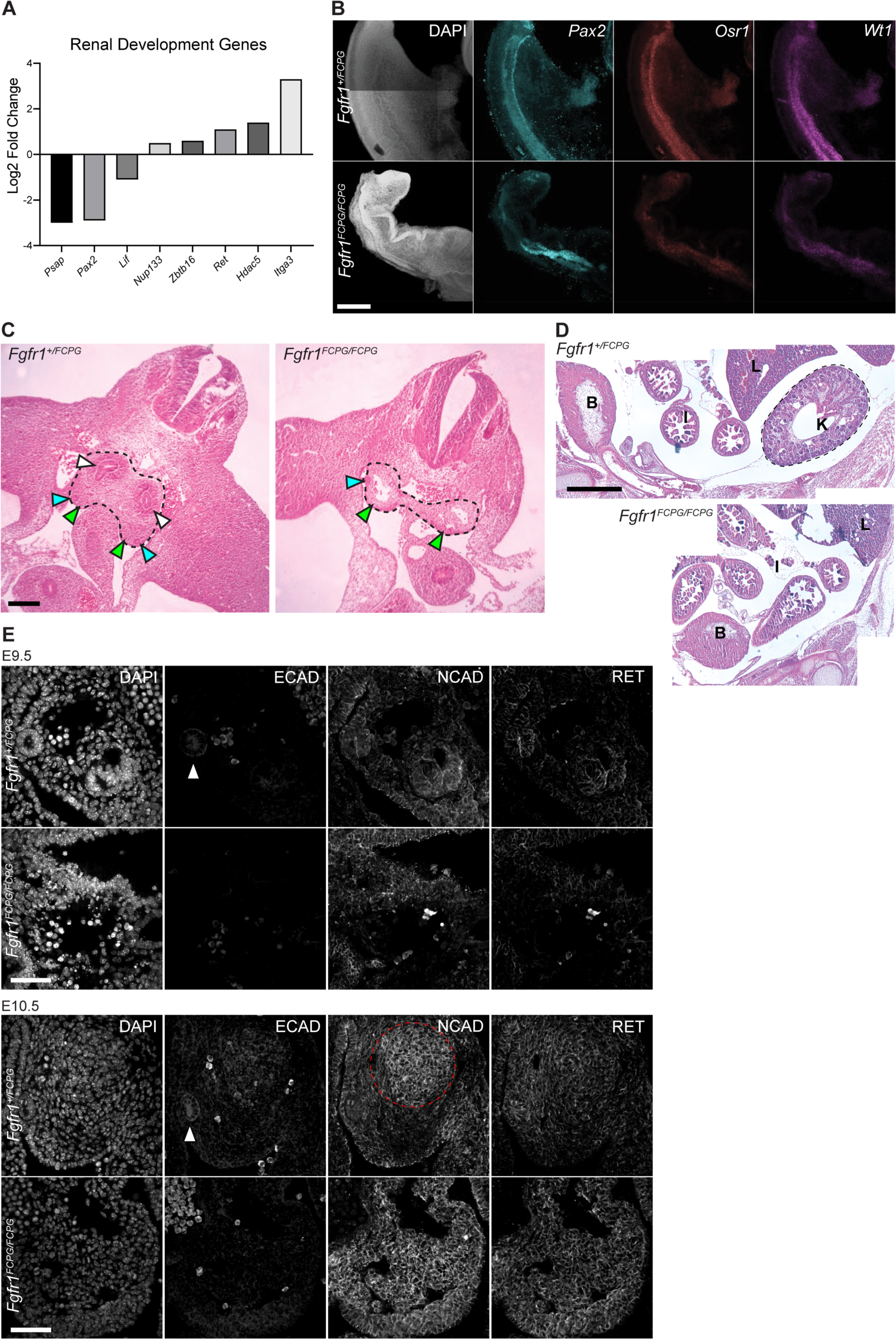
*Fgfr1^FCPG^*mutant embryos exhibit delayed intermediate mesoderm development resulting in kidney agenesis. **A)** RNA-seq analysis of *Fgfr1^FCPG/FCPG^* embryos shows an alteration in expression of renal development genes, including *Pax2* and *Ret*, compared to heterozygotes. **B)** The expression domain of key intermediate mesoderm markers, *Pax2, Osr1,* and *Wt1*, is altered and overall expression levels are reduced in *Fgfr1^FCPG/FCPG^* embryos, compared to controls, at E9.5. Scale bar represents 200μm. **C)** At E11.5, *Fgfr1^FCPG/FCPG^* embryos lack a ureteric bud and the mesonephric mesenchyme does not appear to be condensing, a key process for the early formation of the kidney. Dashed lines encompass the developing intermediate mesoderm. White arrows indicate ureteric buds, blue arrows indicate nephrogenic duct, and yellow areas indicate urogenital ridge. Scale bar represents 10μm. **D)** At E14.5, *Fgfr1^FCPG/FCPG^*embryos lack a recognizable kidney. Wild-type embryos exhibit a distinctive kidney (K) proximal to the caudal body wall below the liver (L). The bladder (B) and intestine (I) are also labeled. Scale bar represents 500μm. **E)** Coordination of Cadherin localization is disrupted in *Fgfr1^FCPG/FCPG^* embryos, at both E9.5 and E10.5, during early mesonephric mesenchyme condensation and ureteric bud formation. ECAD is greatly reduced in the mutants, while NCAD is more broadly localized to the gonadal ridge rather than restricted to the condensing mesonephric mesenchyme. Localization of RET, a key driver of early kidney development, appears to be unaffected in homozygous *Fgfr1^FCPG^* mutants, although expression is slightly higher as also seen in the RNA-seq. White arrows indicate the nephrogenic duct. Dashed red line encompasses the condensing mesonephric mesenchyme. Scale bar represents 50μm.

To identify how intermediate mesoderm was being affected during early development, we used *in situ* analysis to observe key gene expression during development. In addition to *Pax2*, we probed two genes involved in the earliest stages of kidney development, *Osr1* and *Wt1*. While *Pax2* was expressed in *Fgfr1^FCPG^* embryos, its expression was expanded, with a wider range anteriorly and absent expression posteriorly (Fig. 4B). In addition, we saw a reduction in expression levels of both *Osr1* and *Wt1* across both anteriorly and posteriorly in *Fgfr1^FCPG/FCPG^* embryos at E9.5 (Fig. 4B). Together, these data implicate *Fgfr1* at the earliest stages of renal development, playing an earlier role in patterning the intermediate mesoderm and initiating nephric duct formation than previously reported.

Sectioning through the posterior truncation in homozygous *Fgfr1^FCPG/FCPG^*embryos revealed severe defects in kidney formation. At E11.5, heterozygous *Fgfr1^+/FCPG^* embryos showed formation of the ureteric bud, the structure that will give rise to the kidney, while *Fgfr1^FCPG/FCPG^*embryos exhibited a complete lack of ureteric bud formation, did not show signs of metanephric mesenchyme condensation, and exhibited hypoplasia of the gonadal ridge (Fig. 4C). By E14.5, wild-type embryos developed a kidney adjacent to the rostral body wall below the liver. In contrast, at E14.5, *Fgfr1^FCPG/FCPG^* embryos displayed complete agenesis of the kidney (Fig. 4D). Conditional deletion of *Fgf8* using *TCre* resulted in increased cell death in the metanephric mesenchyme that led to kidney hypoplasia (Perantoni et al., 2005) While we did observe altered cell death in the gonadal ridge of *Fgfr1^FCPG/FCPG^*embryos (Fig. S4A), suppression of cell death via *Bim+/-* heterozygosity resulted in no significant change in embryonic development (Fig. S4B), suggesting that reducing cell death is not sufficient to rescue development at E10.5. These phenotypes reveal a more severe effect on kidney development due to loss of canonical *Fgfr1* signaling than previously observed in isoform or conditional mutants (Sims-Lucas et al., 2011). Previously, we had observed defects in bifurcation of the kidneys in *Fgfr1^1′Frs/1′Frs^* mutants, which lack the ability to recruit *Frs2/3* to *Fgfr1*. In these hypomorphic mutants, a single kidney was formed with branched nephrons (Hoch and Soriano, 2006). The more severe kidney agenesis found in *Fgfr1^FCPG/FCPG^*mutants indicates a sufficiency for *Frs2/3*-independent signaling at the earliest stages of kidney development but a necessity for *Frs2/3*-dependent signaling during bifurcation. Interestingly, *Fgfr1^FCPG/FCPG^* mutant embryos still developed gonads, which are also derived from the intermediate mesoderm. This indicates that the pronephros and mesonephros are potentially unaffected, but metanephros development is impaired.

During early renal morphogenesis, a mesenchymal to epithelial transition is required to properly form the ureteric bud. This event requires a coordinated transition from the expression of *Cdh2 (N-cadherin)* to *Cdh1 (E-cadherin)*. To determine if this process is disrupted, we examined NCAD and ECAD protein expression and localization, as well as expression of RET, a key regulator of early kidney development. At E9.5, NCAD was the predominantly expressed cadherin in the intermediate mesoderm, with ECAD being mostly restricted to the Wolffian/Mullerian duct in wild-type embryos (Fig. 4E). *Fgfr1^FCPG/FCPG^* embryos expressed NCAD in the intermediate mesoderm but lacked ECAD expression in the Wolffian/Mullerian duct. RET expression was low in both genotypes at this stage. By E10.5, the differences between *Fgfr1^FCPG/FCPG^*mutant and wild-type embryos were starker. Wild-type mesonephric mesenchyme began to express ECAD at the center of the condensing mesenchyme alongside higher levels of NCAD, while *Fgfr1^FCPG/FCPG^* mutants exhibited very little ECAD and high levels of NCAD across the entire region rather than localized to the mesonephric mesenchyme. RET expression was increased in both genotypes at this stage and did not appear to be perturbed in the *Fgfr1^FCPG/FCPG^*mutants (Fig. 4E). Together, these data indicate that *Fgfr1* plays a major role in early intermediate mesoderm formation at the earliest stages of kidney development.

### A novel Fgfr1^FCSPG^ allele further reduces Fgfr1 activity in the absence of canonical FGF signaling

Because *Fgfr1^FCPG/FCPG^* mutants survive much later than *Fgfr1^-/-^* null mutants on the same genetic background, despite eliminating all known canonical signaling (Brewer et al, 2015, Ray et al. 2020), we introduced a tyrosine to phenylalanine conversion at Y730, which we refer to as S because Y730 has been found to interact with Src family kinases (SFKs), as well as PIK3R2, the p85 regulatory subunit of PI3K (Dudka et al., 2010; Francavilla et al., 2013)(Fig. 5A). Using gene targeting, we constructed two lines, *Fgfr1^SMyc^* carrying the Y730F mutation on its own and *Fgfr1^FCSPGMyc^* carrying Y730F in conjunction with our previous signaling FCPG mutations, both with a 2xMyc tag at the C-terminus (Fig. 5A, Fig. S5A, B). These lines are hereafter referred to as *Fgfr1^S^* and *Fgfr1^FCSPG^*, respectively, for simplicity.

**Figure 5.**
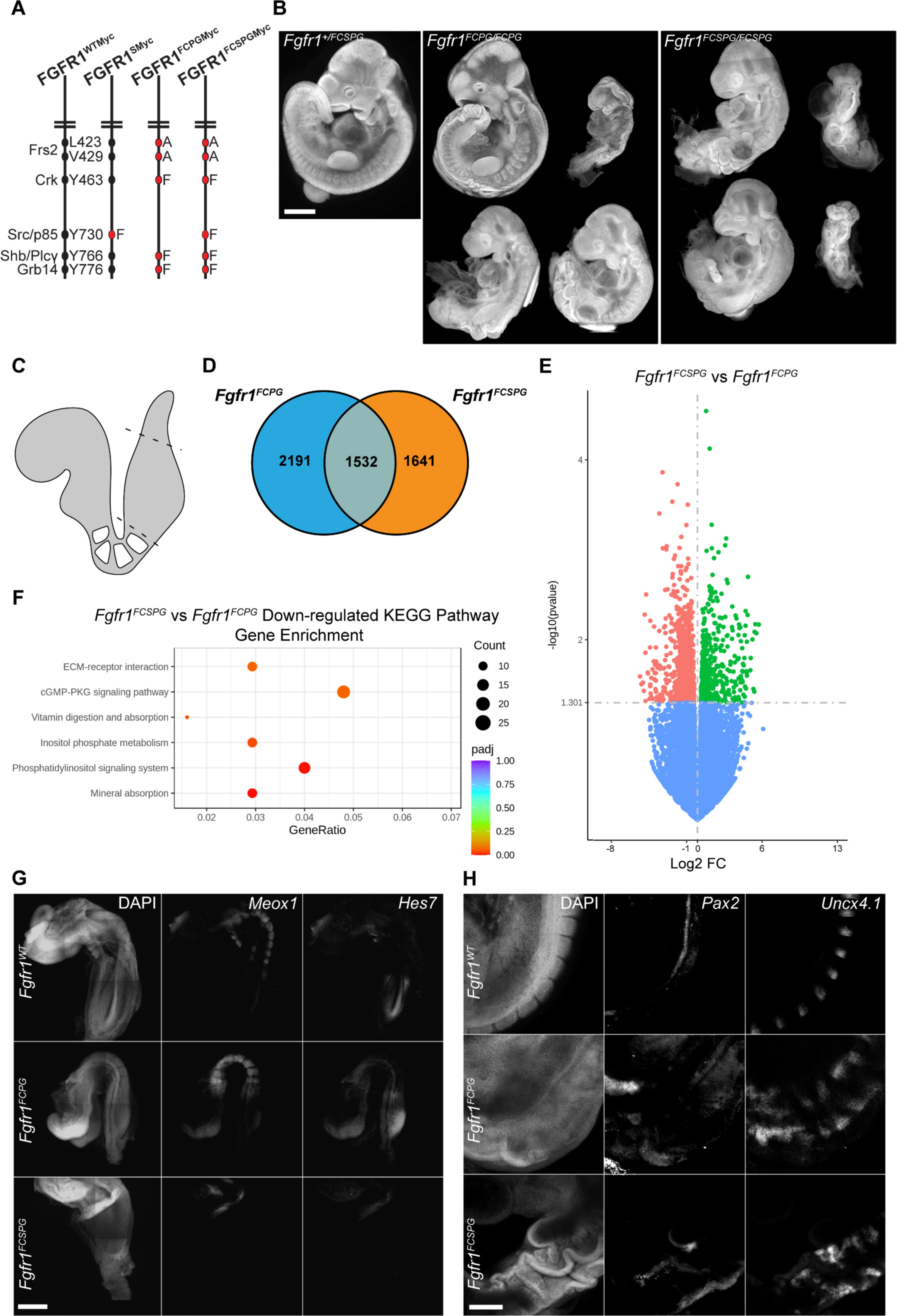
RNA-seq analysis of *Fgfr1^FCPG^* and *Fgfr1^FCSPG^* E8.5 tailbud tissue reveals downregulation of genes involved in ECM interaction and PIP signaling pathways. **A)** Schematic of signaling mutant alleles constructed in *Fgfr1* gene. The most severe allele, *Fgfr1^FCSPGMyc^*, has an additional mutation at Y730, a potential binding site for SFKs and/or p85, in addition to the FCPG mutations that disrupt downstream canonical signaling. **B)** Both *Fgfr1^FCPG/FCPG^* and *Fgfr1^FCSPG/FCSPG^* embryos show severe defects at E10.5 Phenotypes are variable, but most *Fgfr1^FCPG/FCPG^* embryos fail between E9.5 and E10.5, while all *Fgfr1^FCSPG/FCSPG^*fail at or before E9.5. Phenotypes include axial truncations, neural tube closure defects, somitogenesis defects, and cardiac defects. Scale bar represents 1mm. **C)** Tailbud tissue was dissected from wild-type and homozygous *Fgfr1^FCPG^* and *Fgfr1^FCSPG^* embryos at E8.5 to examine the onset of the observed phenotypic changes. Allantois tissue was removed from the tailbud tissue (dashed lines). **D)** Venn diagram depicting significant differentially expressed genes in homozygous *Fgfr1^FCPG^* or *Fgfr1^FCSPG^*mutant tailbud tissue compared to wild-type tissue at E8.5. Volcano plots displaying differential gene expression between *Fgfr1^FCPG/FCPG^* mutants and wild type, *Fgfr1^FCSPG/FCSPG^* mutants and wild type, and *Fgfr1^FCSPG/FCSPG^*mutants and *Fgfr1^FCPG/FCPG^* mutants, respectively, at E8.5. Horizontal dashed line signifies significance cut-off of p=0.05. **F)** KEGG Pathway GSEA of genes downregulated in *Fgfr1^FCSPG^* tissue compared to *Fgfr1^FCPG^* tissue. The most significantly downregulated BP terms revolve around tissue migration, indicating that S mutation may differentially affect *Fgfr1’s* role in cell migration. Additionally, KEGG Pathway GSEA identified ECM-receptor interaction, inositol signaling, and pathways related to calcium regulation. **G)** At E8.5, *Fgfr1^FCSPG/FCSPG^*embryos display a significant reduction in somitogenesis as seen by Meox1 and Hes7 expression. In contrast, *Fgfr1^FCPG/FCPG^* exhibits normal somitogenesis at this stage. Scale bar represents 200μm. **H)** At E9.5, both Fgfr1^FCPG/FCPG^ and *Fgfr1^FCSPG/FCSPG^* exhibit defective patterning in intermediate mesoderm and somitic mesoderm, as marked by *Pax2* and *Uncx4.1*, respectively. Scale bar represents 200μm.

*Fgfr1^S^* mice did not exhibit apparent phenotypes and could be maintained as homozygous breeding stocks. We next shifted our attention to the expectedly more severe combinatorial signaling allele, *Fgfr1^FCSPG^*. As the majority of *Fgfr1^FCPG^* embryos arrest at E10.5, we decided to examine if the *Fgfr1^FCSPG/FCSPG^* embryos exhibited a similar lethality. *Fgfr1^FCSPG/FCSPG^* embryos were recoverable up to E10.5 (Table 1). There was a large degree of variation in the severity of the phenotypes observed, despite the fact that the mutation was analyzed on a co-isogenic genetic background. Mutant embryos exhibited multiple phenotypes including defects in axial extension, neural tube closure, somite patterning, cardiac development, and limb bud development, with the majority appearing to have arrested during turning prior to E9.5 (Fig. 5B). Additionally, while 13/16 *Fgfr1^FCSPG/FCSPG^* embryos had beating hearts at E9.5, only 2/9 did so at E10.5, suggesting that the mutants were necrotic by that stage. Using a qualitative scale based on observed phenotypes (Table 4), we graded the severity of mutants observed for both alleles and found that *Fgfr1^FCSPG/FCSPG^* embryos trended toward more severe than *Fgfr1^FCPG/FCPG^* embryos when collected at E10.5 (Fig. S5C).

### Fgfr1^FCSPG/FCSPG^ mutants display more severe changes in expression of genes involved in migration, inositol signaling, and cell polarity compared to Fgfr1^FCPG//FCPG^ mutants

Due to the earlier lethality of the *Fgfr1^FCSPG/FCSPG^* embryos, we re-analyzed the scRNA-seq dataset examining gastrulation and early organogenesis, revealing multiple peaks of *Fgfr1* expression at E7, E7.75, and E8.5 (Pijuan-Sala et al., 2019)(Fig. S5D). The peak of *Fgfr1* expression at E8.5 aligned with the onset of phenotypes we observed in both *Fgfr1^FCPG/FCPG^* and *Fgfr1^FCSPG/FCSPG^*embryos, indicating a potential necessity for *Fgfr1* activity at this stage. To determine how *Fgfr1* is impacting the earliest stages of intermediate mesoderm development and the increased severity of *Fgfr1^FCSPG/FCSPG^*mutant phenotypes, we performed RNA-seq on E8.5 tailbud tissue. Tailbuds were excised just after the last definitive somite and the allantois was removed to isolate the NMPs (Fig. 5C). Both homozygous *Fgfr1^FCPG/FCPG^* and *Fgfr1^FCSPG/FCSPG^* tailbuds showed a large number of significantly altered genes, with 2191 and 1641 uniquely significant DEGs, respectively, and 1532 common DEGs, compared to wild-type samples (Fig. 5D). We found many common GO BP terms altered in both *Fgfr1^FCPG^* and *Fgfr1^FCSPG^* embryos involving pattern specification and skeletal, limb, and renal development.

As *Fgfr1^FCSPG/FCSPG^*mutants presented more severe phenotypes than *Fgfr1^FCPG/FCPG^* mutants, we next focused on the differential gene expression between the two to identify which pathways were more severely affected. Compared to wild-type embryos, *Fgfr1^FCPG/FCPG^* embryos exhibited 2142 up and 1581 downregulated DEGs. Meanwhile, *Fgfr1^FCSPG/FCSPG^* embryos had 1820 up and 1353 downregulated DEGs compared to wildtype (Fig. S5E). Comparing *Fgfr1^FCSPG/FCSPG^* mutants to *Fgfr1^FCPG/FCPG^* mutants there were 932 upregulated and 550 downregulated DEGs (Fig. 5E, Table 5). GO term GSEA identified downregulation of two groups of BP terms involving cell migration and cell junction assembly, as well as CC terms involving the apical cell membrane (Fig. S6A, B). While we did observe changes in cell migration genes in *Fgfr1^FCPG/FCPG^* embryos, these data suggest that the changes in *Fgfr1^FCSPG/FCSPG^* mutants were more severe. GO Terms involving both cell migration and cell junctions were more significantly impacted in *Fgfr1^FCSPG/FCSPG^* vs *Fgfr1^+/+^* compared to *Fgfr1^FCPG/FCPG^* vs *Fgfr1^+/+^*. Changes in genes localized to the apical membrane could indicate defects in cell polarity. KEGG Pathway GSEA identified phosphatidylinositol signaling (5-fold enrichment, p<0.0001) and ECM-receptor interactions (2.9-fold enrichment, p=0.001) as two of the most significantly altered pathways in *Fgfr1^FCSPG/FCSPG^* mutants (Fig. 5F, Table 6). Phosphatidylinositol signaling is a multifunctional pathway that impacts a wide array of cellular functions, ranging from cell membrane structure to signal transduction, vesicle trafficking, cytoskeletal dynamics, growth, and survival (Anderson et al., 1999; Marat and Haucke, 2016; Wen et al., 2023). DEGs identified include PI4P modifiers as well as genes involved in endocytosis and membrane trafficking. ECM-receptor interaction was identified with significant changes in multiple integrins and ECM components. We queried many of the affected genes found under these KEGG Pathway terms against the scRNA-seq databases (Cao et al., 2019; Pijuan-Sala et al., 2019) and found many showed overlapping expression with *Fgfr1* at E8.5 (Fig. S7, S9) and from E9.5 to E11.5 (Fig. S8, S10).

### Fgfr1^FCSPG/FCSPG^ embryos show earlier onset of disruptions to somitic mesoderm compared to Fgfr1^FCPG^ embryos

RNA-seq analysis of *Fgfr1^FCPG/FCPG^* and *Fgfr1^FCSPG/FCSPG^*mutants indicates that both *Meox1* and *Pax2*, key markers of somitic and intermediated mesoderm respectively, are downregulated at E8.5. As the most severe *Fgfr1^FCSPG/FCSPG^* embryos collected at E10.5 appear to have arrested between E8.5 and E9.5 (Fig. 5B), we opted to examine somitic and intermediate mesoderm through *in situ* analysis at both stages. Somitic mesoderm appeared to be disrupted as early as E8.5 in *Fgfr1^FCSPG/FCSPG^* embryos, which exhibited a lack of defined somites, as evidenced by the loss of *Meox1*, and disruption to the somitogenesis clock, as evidenced by the absence of *Hes7* in the tailbud (Fig. 5G). Similar to *Fgfr1^FCPG/FCPG^* mutants, homozygous *Fgfr1^FCSPG/FCSPG^* embryos displayed altered intermediate and somitic mesoderm patterning at E9.5, as marked by *Pax2* and *Uncx4.1*, respectively (Fig. 5H).

FGF signaling is a key regulator of the somitogenesis clock, along with Wnt and Notch (Wahl et al., 2007). To determine if the three pathways were further disrupted, we dissected the tailbuds of E9.5 embryos, extracted mRNA, and analyzed gene expression via qRT-PCR. Interestingly we did not see a change in downstream FGF targets, however we did see downregulation of *Dkk1*, a Wnt negative regulator, and *Hes7* and *Nrarp*, two Notch targets (Fig. S5F). There is known crosstalk between FGF, Wnt, and Notch signaling during somitogenesis, with each pathway engaging feedback loops that regulate both themselves and the other two pathways (Déqueant and Pourquié, 2008). The *in situ* data indicates that the *Fgfr1^FCSPG^* allele has a greater impact on somitogenesis than the *Fgfr1^FCPG^* allele, as *Hes7* expression is absent in *Fgfr1^FCSPG/FCSPG^*embryos. As the *Fgfr1^FCPG^* allele eliminates all canonical FGF signaling, this suggests the presence of additional mechanisms beyond canonical signaling exist that regulate mesodermal development and somitogenesis.

### FGFR1 alleles exhibit differing endocytic trafficking patterns

As *Fgfr1^FCSPG/FCSPG^* mutants exhibited a more severe reduction in mesodermal development compared to *Fgfr1^FCPG/FCPG^*mutants, we next wanted to identify what proteins might be recruited to the binding site of Y730 to provide additional function beyond canonical FGF signaling. To determine what proteins were binding to FGFR1, we performed a co-immunoprecipitation (Co-IP) using FGFR1^WT-3xFlag^, FGFR1^FCPG-3xFlag^, and FGFR1^FCSPG-3xFlag^ proteins expressed in NIH3T3 fibroblasts. Using label-free quantification mass spectroscopy (MS), we identified proteins that were present in two different scenarios. First, we identified proteins that would still bind to all three FGFR1 forms to reveal interactions that might provide insight into what function FGFR1 maintains, even in the most severe FGFR1^FCSPG^ mutant protein. Second, we identified proteins that are bound to both Fgfr1^WT^ and FGFR1^FCPG^ but not FGFR1^FCSPG^. These proteins would potentially indicate what function Y730 was providing to FGFR1, and what could account for the difference in phenotypes observed during development.

Across all six samples, *Fgfr1^WT-3xFlag^, Fgfr1^FCPG-3xFlag^,* and *Fgfr1^FCSPG-3xFlag^* with or without FGF1 treatment, 1418 proteins were identified after removing background signal. GO Term analysis classified 549 of these proteins (31.4%) as involved in cellular processes, followed by metabolic processes (17.3%), biological regulation (13.1%), and localization (7.6%) (Fig. 6A, left panel). We focused on the proteins involved in cellular processes, as defined by any process that acts at the cellular level including cell communication, cell growth, cell division, and signal transduction, and further divided them into secondary and tertiary terms. As both ECM-receptor interactions and PIP signaling were identified in the RNA-seq, we focused on GO BP Terms involving either of these processes. We additionally highlighted several terms involving endocytosis, as PIP signaling plays a role in regulating endosomal trafficking. Of those GO BP Terms, some of the most enriched terms were Clathrin-dependent endocytosis (5.8-fold enrichment, p<0.0001) and regulation of focal adhesion assembly (4-fold enrichment, p<0.0001) (Fig. 6A, right panel, Table 7).

**Figure 6.**
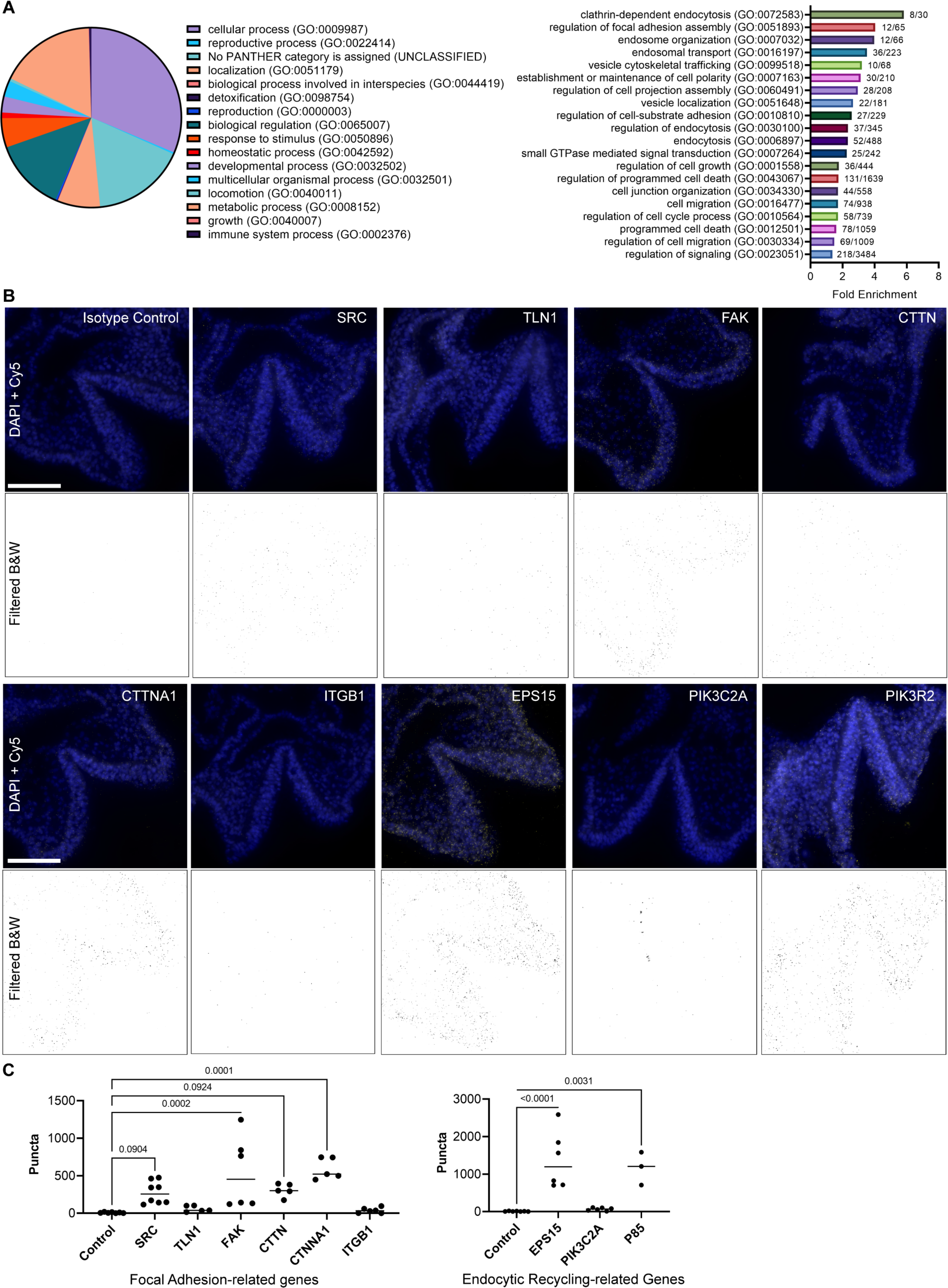
Protein analysis of FGFR1^FCPG^ and FGFR1^FCSPG^ reveals FGF receptor interaction with focal adhesion complexes and intracellular trafficking. **A)** GO Term analysis of proteins identified from Co-immunoprecipitation/Mass spectroscopy. Top-level primary term distribution from 1418 proteins identified across all FGFR1 mutants and control (left panel). Fold-enrichment of secondary and tertiary terms involved in endocytosis and ECM interactions. **B)** Images of proximity ligation assay in E8.5 *Fgfr1^Myc/Myc^*embryos. An anti-Myc antibody was paired with an antibody against the indicated protein to identify proximity of the two proteins within 40nm. Background fluorescence was filtered, and threshold values were standardized across all images. Scale bar represents 100μm. **C)** Quantification of PLA puncta in focal adhesion-related genes (left graph) and endocytosis-related genes (right graph). FAK, CTNNA1, EPS15, and PIK3R2 all exhibited a significant increase in puncta compared to control samples. SRC and CTTN displayed an increased trend in puncta but were not statistically significant. Significance was determined using One-way ANOVA with Bonferroni correction for multiple comparisons.

From the Co-IP MS we found that the SFK FYN associated with FGFR1^WT-3xFlag^, FGFR1^FCPG-3xFlag^, and FGFR1^FCSPG-3xFlag^ receptors, indicating that it was still recruited even with the S mutation, possibly through a mechanism not involving the conserved SFK SH2 domain since other SFKs did not associate with FGFR1. Alongside FYN was Cortactin (CTTN), Catenin-alpha (CTNNA1), and Talin-1 (TLN1), all components of focal adhesion complexes involved in integrin-mediated cell adhesion and regulatory targets of SFKs (Mitra and Schlaepfer, 2006; Ortiz et al., 2021). To examine the validity of these interactions *in vivo*, we performed PLA in E8.5 *Fgfr1^Myc/Myc^* embryos. Using available antibodies, we probed endocytic FGFR1 interactions with SRC, TLN1, Focal adhesion kinase (FAK), CTTN, CTNNA1, and Integrin beta 1 (ITGB1). Quantification revealed significant interactions with FAK and CTNNA1 (p<0.001), indicating proximity with FGFR1. SRC and CTTN exhibited increased puncta compared to controls but did not reach statistical significance (p=0.09); the SRC antibody cross-reacts with FYN and YES indicating potential interaction with any of these three broadly expressed SFKs. TLN1 and ITGB1 showed little to no interaction with FGFR1 (Fig. 6B, C). The interaction with both FAK and CTNNA1 indicates that FGFR1 interacts closely with proteins involved in focal adhesion assembly and may play a role in these structures.

Interestingly, multiple proteins involved in intracellular trafficking were found in the FGF-stimulated FGFR1^WT^ and FGFR1^FCPG^, but not the FGFR1^FCSPG^, Co-IP MS samples. Vesicle-associated membrane protein B (VAPB) and Epidermal growth factor receptor substrate-15 (EPS15) are two proteins involved in intracellular membrane trafficking and endocytic recycling. Kinesin family member-2a (KIF2A), Microtubule-associated protein-6 (MAP6), and Cytoskeleton-associated protein-5 (CKAP5) are proteins involved with microtubule-mediated transport. Together these interactions indicate that Y730 may be playing a role in mediating the endocytic trafficking of FGFR1, much like the congruent Y734 residue in FGFR2.

We again used PLA to examine interactions with proteins involved in endocytic trafficking: EPS15, PIK3C2A, and PIK3R2 (p85). Both EPS15 and PIK3R2 displayed significantly increased puncta compared to controls (p<0.0001 and p=0.0031, respectively), while PIK3C2A showed no significant difference (Fig. 6B, C). PIK3R2 interaction with FGFR1 had previously been shown during induced endocytic recycling of the receptor (Francavilla et al., 2013). Additionally, EPS15 is a known component of clathrin-coated pits and plays a role in the endocytic regulation of EGFR and potentially other RTKs (Haugen et al., 2017; Salcini et al., 1999). The combined Co-IP MS and PLA data suggest that the S mutation (Y730F) inhibits association with endocytic regulatory proteins.

To determine whether endocytic trafficking was substantially affected *in vivo,* we examined FGFR1 colocalization with EEA1, a marker of early endosomes, and RAB7, a marker for mid to late endosomes. Using super-resolution stimulated emission depletion (STED) microscopy we observed colocalization in wild-type, *Fgfr1^FCPG/FCPG^*, and *Fgfr1^FCSPG/FCSPG^* embryos in E9.5 somitic mesoderm. FGFR1^FCPG^ exhibited an increase in association with early endosomes compared to FGFR1^FCSPG^, with both showing significant increases over FGFR1^WT^ (Fig. 7A). When looking at later stage endosomes, FGFR1^FCPG^ again displayed the greatest association compared to both FGFR1^WT^ and FGFR1^FCSPG^, which exhibited similar association rates compared to each other (Fig. 7B). These data indicate that the mutant FGFR1 proteins are trafficked differently than wild-type FGFR1. Interestingly, the data also suggest that FGFR1^FCPG^ and FGFR1^FCSPG^ are trafficked differently compared to each other (Fig. 7C). Together, the Co-IP, PLA, and STED microscopy data support previous data suggesting that FGFR1 Y730 may be involved in regulating endocytosis of the receptor. These changes in endocytic sorting may account for the reduced functionality of the *Fgfr1^FCSPG^* allele, resulting in the more severe phenotypes observed in *Fgfr1^FCSPG/FCSPG^* embryos compared to *Fgfr1^FCPG/FCPG^*.

**Figure 7.**
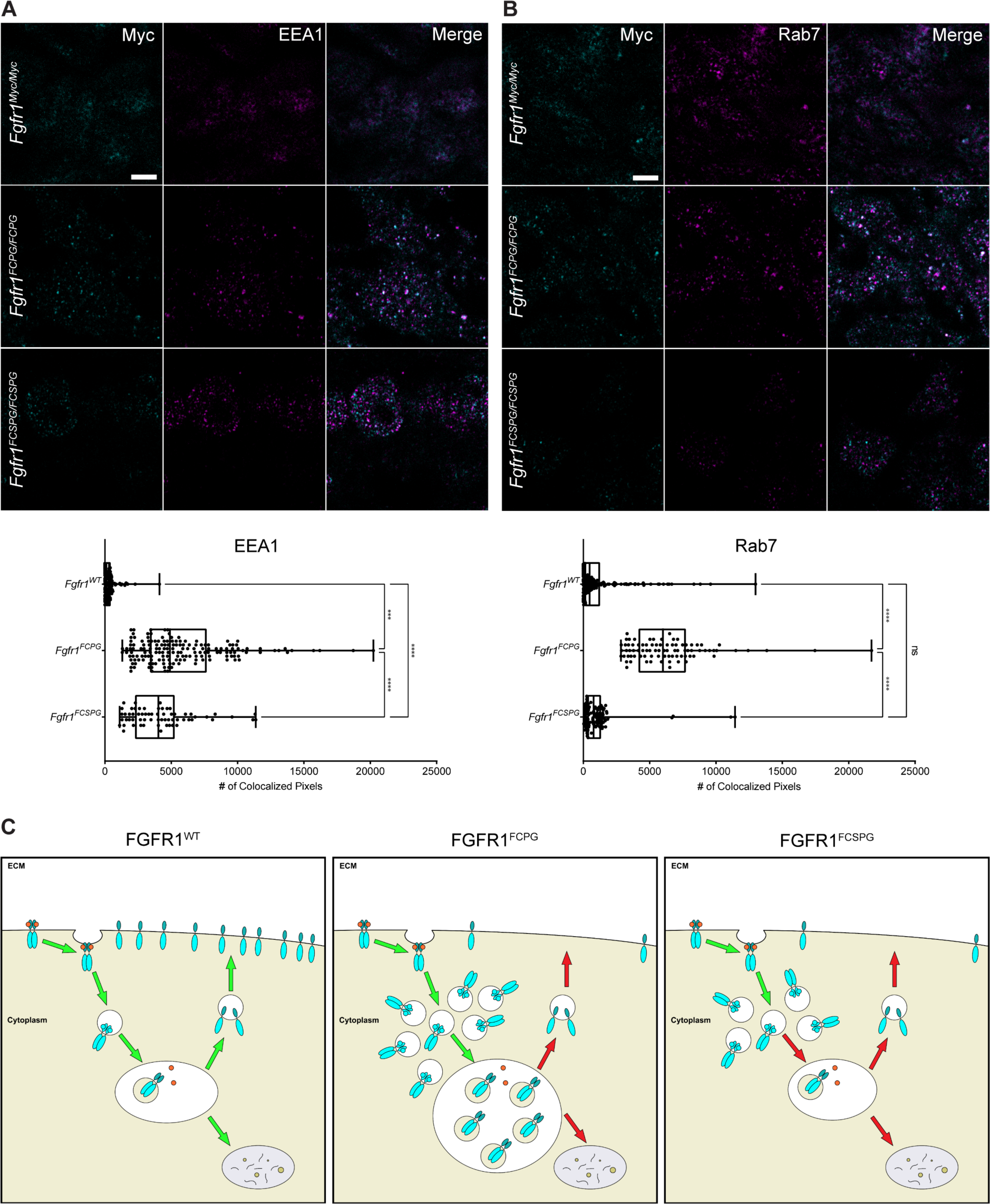
FGFR1^FCPG^ and FGFR1^FCSPG^ exhibit alterations in endocytic trafficking and recycling. **A)** Colocalization of FGFR1 with EEA1, a marker of early endosomes. Both FGFR1^FCPG^ and FGFR1^FCSPG^ show an increase colocalization, with FGFR1^FCPG^ exhibiting significantly more colocalization than either FGFR1^FCSPG^ or FGFR1^Myc^. Significance was determined using One-way ANOVA with Bonferroni correction for multiple comparisons. Scale bar represents 3μm. **B)** Colocalization of FGFR1 with RAB7, a marker of mid to late endosomes. FGFR1^FCPG^ exhibits a significant increase in colocalization compared to either FGFR1^Myc^ or FGFR1^FCSPG^. Significance was determined using One-way ANOVA with Bonferroni correction for multiple comparisons. Scale bar represents 3μm. **C)** Proper endocytic recycling of FGFR1 is required to maintain full functionality of FGFR1 signaling. FGFR1^FCPG^ accumulates in EEA1 and Rab7-positive endosomes, indicating a reduction in recycling to the membrane. FGFR1^FCSPG^ also exhibits changes in trafficking with accumulation in EEA1-positive endosomes, indicating a failure to proceed through endocytic recycling pathways.

## Discussion

Previous studies of both *Fgfr1* and *Fgfr2* allelic series of mutants have shown that both receptors exhibit activity outside of canonical FGF signaling pathways (Brewer et al., 2015; Ray et al., 2020; Ray and Soriano, 2023). We observed that some *Fgfr1^FCPG/FCPG^* embryos, devoid of canonical signaling, die perinatally. Further analysis revealed severe defects in somitic, intermediate, and lateral plate mesoderm. Surviving embryos had severe axial truncations, with somitic mesoderm defects beginning at E8.5. We observed complete kidney agenesis with defects in intermediate mesoderm organization at E9.5. Additionally, digit patterning defects were observed, and ossification of bone was delayed throughout later development. To further dissect the remaining function in *Fgfr1* we introduced an additional point mutation at Y730 and generated *Fgfr1^FCSPG^*mutants. Using a combination of RNA-seq and protein interaction analyses, we found that FGFR1 interacts with proteins involved in focal adhesions and endocytic trafficking. Accordingly, using super-resolution microscopy, we identified alterations in endocytic trafficking between wild-type and mutant FGFR1. Through this work, we establish the importance of non-canonical FGF signaling in mesoderm development and highlight the role of endocytic trafficking in FGF receptor signaling.

Recovery of *Fgfr1^FCPG/FCPG^* embryos at P0 highlights the surprising observation that canonical *Fgfr1* signaling is dispensable for many developmental processes. The discrepancy between our current data and the previous, smaller scale *Fgfr1^FCPG^* recovery data (Brewer et al., 2015) could be due to two possibilities. First, we maintained the new *Fgfr1^FCPGMyc^* line by crossing heterozygous males and females, allowing us to potentially recover homozygous animals that survive until birth, even at low frequencies. During the initial analysis of the *Fgfr1^FCPG^*allele, the colony was maintained as heterozygous males crossed to wild-type females, so large numbers of homozygous animals would never have been generated in the first place to detect rare embryos that survived until birth. Alternatively, there may have been genetic drift in the 129S4 background over the years, resulting in reduced sensitivity to partial loss of *Fgfr1* function.

While the posterior skeletal anatomy is affected in *Fgfr1^FCPG/FCPG^*embryos, anterior anatomy is comparatively normal. Gross observations of the heart, lungs, and digestive system revealed no apparent phenotypes. The most severely affected processes observed in these mutants were ossification, digit patterning, posterior axial extension, and kidney development. Since both posterior axis formation and early kidney development occur concurrently, these data highlight a major role for *Fgfr1* signaling around E8.5. As only one third of *Fgfr1^FCPG/FCPG^*, and no *Fgfr1^FCSPG/FCSPG^*, embryos develop past E10.5, we hypothesize that a minimal level of *Fgfr1* activity is required during early mesoderm differentiation to continue development. The signaling mutant alleles that do not provide sufficient activity result in halted development, most likely due to insufficient mesoderm expansion. However, the *Fgfr1^FCPG^* allele may be straddling that threshold. Minor changes in cellular environments or stochastic events may allow some *Fgfr1^FCPG/FCPG^* embryos to breach that minimal level of activity, providing just enough mesoderm to continue development, but not to produce all mesoderm-derived structures.

Both *Fgfr1^FCPG/FCPG^* and *Fgfr1^FCSPG/FCSPG^* mutants display deficiencies in somitic, intermediate, and lateral plate mesoderm. Surprisingly, both mutants form anterior somites, indicating that canonical FGFR1 signaling is not required for initial somite formation. However, somite morphology is affected in both *Fgfr1^FCPG/FCPG^*and *Fgfr1^FCSPG/FCSPG^* embryos indicating that canonical FGFR1 signaling is necessary for proper pacing of the wavefront during somitogenesis. The changes in *Hes7* expression in *Fgfr1^FCSPG/FCSPG^* mutants indicates that this may be the case. Epithelialization of somites during compaction requires the induction of apical basal polarity which would influence changes in ECM interactions and cell-cell adhesion, as observed through RNAseq analysis. Alterations to how *Fgfr1* functions in these processes could result in delayed induction producing improperly spaced and sized somites, as seen in both *Fgfr1^FCPG/FCPG^* and *Fgfr1^FCSPG/FCSPG^* mutants. Alongside depleted somitic mesoderm, lack of intermediate mesoderm development also led to defects in renal development in *Fgfr1^FCPG/FCPG^* mutants. Previous studies have identified *Fgfr2* as a major factor in kidney development, with *Fgfr1* playing a more minor role (Poladia et al., 2006; Sims-Lucas et al., 2011; Zhao et al., 2004). Here we show that canonical signaling from FGFR1 is key to early kidney development, as *Fgfr1^FCPG^* mutants display delayed intermediate mesoderm formation and complete kidney agenesis. However, this defect in the intermediate mesoderm is specific to the development of the ureteric bud, as embryos still produce gonads, indicating a specific impairment of the metanephros. We also did not observe defects in the anterior formation of the mesonephros (data not shown), highlighting the specificity of the defects to the most posterior structures of the intermediate mesoderm. Similarly, posterior lateral plate mesoderm (LPM) development was disrupted in *Fgfr1^FCPG/FCPG^* embryos. Mutant embryos that survived past E10.5 exhibited hypomorphic pelvic bones and both fore and hindlimb defects, all of which develop from the LPM. Together these data indicate that noncanonical FGFR1 signaling is sufficient to initiate NMP differentiation into somitic mesoderm and LPM. However, canonical signaling is required for maintenance of the posterior mesoderm and initiation of kidney development.

*In situ* imaging indicates that cadherin switching is altered in mutants, and RNA-seq analysis identified a significant change in ECM-integrin interactions. FGF is a known regulator of cadherin switching, by regulating *Snai1* expression (Ciruna and Rossant, 2001) during the process of EMT (Gonzalez and Medici, 2014). For the proper formation of the ureteric bud, a MET is required when the intermediate mesoderm begins to epithelialize to form the metanephric duct, which then branches into the surrounding metanephric mesenchyme thus forming the ureteric bud. This process also requires a remodeling of the ECM as the branching occurs. Our previous and current data suggests that FGF signaling is playing a role in both mechanisms. Previous data on salivary gland branching showed that loss of *Fgfr1* and *Fgfr2* resulted in adhesion defects and ruptures to the basement membrane, indicating a role in maintaining ECM interactions (Ray and Soriano, 2023). Our newest data further substantiates this role.

Recent data has suggested that *Fgfr1* may help stabilize cadherins at the cell membrane in conjunction with Src activation (Nguyen et al., 2019). Additionally, the loss of SFKs, Src, Fyn, and Yes, result in changes in localization of FGFR1 indicating an ability for SFKs to regulate the distribution of FGFR1 (Sandilands et al., 2007). Proteomic analysis identified FYN and multiple components of focal adhesions as being able to bind to all FGFR1 variants. These interactions suggest that FGFR1 may play a role in regulating focal adhesion formation through direct association. As the S mutation has not eliminated ECM-adhesion interactions, as observed from the Co-IP analysis, together these data further identify potential interaction with focal adhesion components as an additional means by which FGFR1 is able to function outside of canonical signal transduction. However, there are two possible mechanisms by which FGFR1 may be interacting with focal adhesion components. One mechanism would be through the direct recruitment of FGFR1 to focal adhesions. FGFR1 may bind to both regulatory and structural components of focal adhesions to regulate their turnover, maturity, or longevity to modulate cell migration. Previous evidence identified both FGFR1 and FGFR2 as regulators of focal adhesion turnover during keratinocyte wound repair and in primary neural crest cells (Meyer et al., 2012; Ray et al., 2020). Alternatively, FGFR1 may bind to focal adhesion components outside of focal adhesions. FGFR1 could regulate focal adhesion formation or turnover through association with focal adhesion components in other compartments of the cell, such as endosomes. Lack of proper turnover and recycling of FGFR1 and other focal adhesion components may result in the inability to maintain proper cell migration processes.

One of the key pathways identified in our RNA-seq analysis as being differentially altered between *Fgfr1^FCPG^*and the novel *Fgfr1^FCSPG^* mutants was phosphatidylinositol (PI) signaling. The most prominent connection between FGF signaling and phosphatidylinositol signaling is through PI3K-mediated conversion of PIP_2_ to PIP_3_ and the resulting activation of AKT, however there are a multitude of processes that involve the various products of inositol signaling. Multiple PI species play varied roles in intracellular trafficking including endocytosis, receptor recycling, multivesicular body sorting, and autophagy. Previous evidence suggested that FGFR2 may engage alternate routes of endocytic processing based on the recruitment of PIK3R2 (Francavilla et al., 2013). Additionally, FGFR2 internalization and trafficking is kinase-dependent (Auciello et al., 2013), indicating that transphosphorylation and recruitment of adaptors is integral to internalization and endocytic trafficking. Our data point to a similar mechanism occurring with FGFR1 activity. While canonical PI3K/AKT signaling is disrupted in *Fgfr1^FCPG^* mutants, as evidenced by a lack of AKT phosphorylation (Ray et al., 2020), RNA-seq analysis of *Fgfr1^FCSPG^* mutants identified a significant reduction in enzymes involved in the production of PI(4,5)P and PI(3,5)P, which play roles in endocytic recycling.

EEA1, a key regulator of early endosome trafficking, directly binds to PI_3_P, tethering it in the endosomal membrane to aid in regulating early endosome fusion and sorting (Simonsen et al., 1998). We observed increased colocalization of both FGFR1^FCPG^ and FGFR1^FCSPG^ with EEA1, compared to wild-type FGFR1, in somitic mesoderm. Additionally, we observed an increase in colocalization between FGFR1^FCPG^ and RAB7, a regulator of endosome maturation. These data indicate that all three FGFR1 proteins are behaving differently during endosomal trafficking. During normal somitic development, it appears that wild-type FGFR1 is broadly localized across the cell membrane, with less localization to either early or late endosomes. This indicates that FGFR1 is predominantly recycled to the cell membrane, enabling the mesodermal cells to readily respond to incoming FGFs. However, both FGFR1^FCPG^ and FGFR1^FCSPG^ appear to behave differently. Overall FGFR1^FCPG^ localization appears to be mainly in internal puncta, unlike wild-type FGFR1. Changes in FGFR1^FCPG^ localization were expected, as interaction with canonical effectors, such as PLCγ, are known to impact receptor internalization (Sorokin et al., 1994). While loss of PLCγ interaction alone results in reduced internalization, loss of interaction with other canonical effectors in the *Fgfr1^FCPG/FCPG^*mutants may further alter the internalization and localization of FGFR1^FCPG^. Consequently, FGFR1^FCPG^ heavily localizes to both early and late endosomes, indicating that the receptor may be predominantly internalized and maintained within the endocytic network, rather than being recycled to the cell membrane. While FGFR1^FCSPG^ also appears to be mainly localized to internal puncta, the distribution is different from either wild-type FGFR1 or FGFR1^FCPG^. FGFR1^FCSPG^ appears to predominantly localize to early endosomes, with little localization to late endosomes. This observation indicates that FGFR1^FCSPG^ may be unable to progress to mature endosomes for proper sorting or recycling. Additionally, the different localization patterns of FGFR1^FCPG^ and FGFR1^FCSPG^ indicates that the S mutation may indeed contribute to the regulation of its internalization and trafficking. As interaction between FGFR2 and PIK3R2 promotes recycling of FGFR2 to the cell membrane, loss of this interaction in FGFR1^FCSPG^ may promote the accumulation of FGFR1 in early endosomes.

Proper trafficking of receptors is key to maintaining sufficient signal response to extracellular stimuli. Previous studies profiling phosphotyrosine activation downstream of FGFRs have identified numerous proteins involved in endocytic trafficking (Auciello et al., 2013; Cunningham et al., 2010; Francavilla et al., 2013; Watson et al., 2022). Multiple trafficking proteins were also identified via a BioID biotinylation assay using *Fgfr1-BirA\**, further supporting a direct interaction between FGFR1 and key endocytic regulatory proteins (Kostas et al., 2018). The increased severity seen in the *Fgfr1^FCSPG/FCSPG^* mutant embryos may indicate that FGFR1^FCSPG^ is unable to be properly transported to the cell membrane when necessary, or may not be recycled properly, reducing the overall ability of the pathway to be engaged. In *Fgfr1^FCPG/FCPG^* embryos, while the receptor is operating without canonical signaling, its endocytic localization is altered from either wild type or *Fgfr1^FCSPG/FSPG^* mutants, possibly allowing for the maintenance of selective function through non-canonical activity and providing enough activity to allow development to continue. As the *Fgfr1^FCSPG/FCSPG^* mutants also display more severe defects in tissue migration and morphology, this disruption to endocytic trafficking may also influence cell adhesion through integrin or cadherin turnover.

Taken together, these data establish the importance of non-canonical FGF signaling in mesoderm development. *Fgfr1^FCPG^* alleles devoid of canonical downstream signals provide sufficient activity to progress through development beyond what was previously expected. Additionally, genetic dissection using the *Fgfr1^FCSPG^*allele highlights a connection between *Fgfr1* and cell adhesion through recruitment to focal adhesions, which may then be modulated through intracellular trafficking and endocytic recycling by PIP_2_ signaling. Further investigations will examine how alterations to these cellular processes result in disruptions to developmental processes during early mesoderm specification and differentiation.

## Acknowledgements

We thank Jia Li, Chantel Dixon, and Leah Naraine for assistance with genotyping and cell culture. We thank Colin Dinsmore, Rob Krauss, and Prash Rangan for discussions and critical comments on the manuscript. We thank Kevin Kelley and the Mouse Transgenic Core for stable cell culture facilities. Microscopy was performed at the Microscopy and Advanced Bioimaging CoRE at the Icahn School of Medicine at Mount Sinai. RNA sequencing of E9.5 to E11.5 tissue was performed at the Genomics CoRE Facility at the Icahn School of Medicine at Mount Sinai. Timely RNA sequencing of E8.5 tissue was performed by Novogene Corporation, Inc.

## Author contributions

P.S. conceptualized the study. J.F.C. and P.S. created the targeting constructs. P.S. performed the gene targeting and blastocyst injections. J.F.C. and P.S performed the investigations. J.F.C. and P.S analyzed the data. J.F.C. wrote the original draft of the manuscript. J.F.C and P.S. reviewed and edited the manuscript. P.S. and J.F.C. acquired the funding. P.S. provided ample artistic advice, entertainment, and sustenance.

## Data Availability

All relevant data can be found within the manuscript and supplementary information. Raw RNAseq datasets can be found at the Gene Expression Omnibus (GEO) under #####. Raw proteomics dataset can be found at the International Molecular Exchange Consortium (IMEx) under ####.

## Materials and Methods

### Animal husbandry

All animal experimentation was conducted according to protocols approved by the Institutional Animal Care and Use Committee of the Icahn School of Medicine at Mount Sinai (LA11-00243). Mice were kept in a dedicated animal vivarium with veterinarian support. They were housed on a 13 hr-11hr light-dark cycle and had access to food and water ad libitum.

### Mouse models

*Fgfr1^tmSor10.1^*, referred to as *Fgfr1^FCPG^*, *Meox2^tm1(cre)Sor^*, referred to as *Meox2-Cre*, and *Bcl2l11^tm1.1Ast^*, referred to as *Bim*, were previously described (Bouillet et al., 1999; Brewer et al., 2015; Tallquist et al., 2000). *Fgfr1^tm13.1Sor^*, referred to as *Fgfr1^WT-Myc^*, *Fgfr1^tm16.1Sor^*, referred to as *Fgfr1^S-Myc^*, *Fgfr1^tm15.1Sor^*, referred to as *Fgfr1^FCPG-Myc^*, *Fgfr1^tm14.1Sor^*, referred to as *Fgfr1^FCSPG-Myc^*, and *Fgfr2^tm9.1Sor^*, referred to as *Fgfr2^WT-Flag^*, were generated by gene targeting. Linearized targeting vectors were electroporated into 129S4 AK7 ES cells. Targeting events were screened by PCR and subsequent restriction digest to detect incorporation of nucleotide substitutions. Proper targeting was verified via Southern blotting using 5’ external, 3’ external, internal, and neo probes. ES cell chimeras were bred to *Meox2-Cre* deleter mice to remove the neomycin selection cassette, and the *Meox2-Cre* allele was subsequently crossed out. All mice were maintained on a 129S4 co-isogenic background.

The *Fgfr1^tm13.1Sor^* and *Fgfr2^tm9.1Sor^* are available from the Jackson Laboratory Repository with stock numbers 039109 and 039110, respectively. The *Fgfr1^tm14.1Sor^* and *Fgfr1^tm15.1Sor^* mice are available from the Mutant Mouse Regional Resources Center (MMRRC) at the Jackson Laboratory with stock numbers 071669 and 071670, respectively.

### Embryological Grading

Embryos were dissected at E10.5, fixed, and stained with DAPI overnight. Morphological defects were graded based on the presence of phenotypes listed in Table 4. Embryos were given one point for each phenotype and one to three points based on overall size of the embryo. As scale was qualitative, we could not perform statistical analysis on the phenotypic scores. Images were taken on a Zeiss Axio Observer microscope fitted with a Hamamatsu Orca-Flash 4.0 LT Plus camera.

### Skeletal preparations

E14.5, E16.5, E17.5, E18.5, and P0 embryos were skinned, eviscerated, and fixed in 95% ethanol overnight. Staining was performed with Alcian blue/Alizarin red (0.015% Alcian blue, 0.005% Alizarin red, 5% glacial acetic acid, 70% ethanol) overnight at 37°C. Skeletons were subsequently cleared in 1% KOH for 24hrs then transferred through a glycerol:KOH series of increasing glycerol concentration and decreasing KOH concentration. Skeletons were photographed in 80% glycerol using a Nikon SMZ-U dissecting scope fitted with a Jenoptik ProgRes C5 camera. Images of full P0 skeletons were merged using the Adobe Photoshop Photomerge function.

### Histology

E10.5, E11.5, and E14.5 embryos were fixed in 4% PFA in PBS at 4°C, rinsed in PBS, dehydrated through an ethanol series, and then embedded in paraffin. Six-micron sections were cut using a Leica microtome, mounted on slides, and stained with Harris modified hematoxylin and Eosin Y using a standard protocol. Images were taken using a Zeiss Axioplan microscope fitted with a Jenoptik ProgRes C5 camera. If needed, images were merged using the Adobe Photoshop Photomerge function.

### Immunohistochemistry and Immunofluorescence

FFPE tissue sections were deparaffinized and rehydrated through a series of decreasing ethanol washes. Heat induced epitope retrieval was performed using a 20mM sodium citrate pH 6.0 solution in a pressure cooker for 30 minutes followed by pressure release and gradual cooling to 45°C. Sections were washed in PBS, blocked in PBS containing 10% goat serum, incubated with primary antibody overnight at 4°C, then with secondary antibody for one hour at room temperature. For IHC, signal was developed using the ImmPACT DAB Substrate Kit (Vector Laboratories SK-4105). Images were taken using a Zeiss Axioplan microscope fitted with a Jenoptik ProgRes C5 camera. For immunofluorescence, images were taken using a Zeiss LSM780 Confocal microscope. For super-resolution STED microscopy, images were taken using a Leica TCS SP8 Confocal microscope. Colocalization was determined using FIJI ImageJ. Threshold values for each channel were calculated for each set of images. Colocalization values were determined using the Colocalization Threshold plugin of ImageJ. One-way ANOVA with Bonferroni correction for multiple comparisons was used to determine statistical significance.

### *In situ* hybridization

Whole mount embryos were fixed in 4% PFA in PBS at 4°C, rinsed in PBS, then dehydrated through a series of methanol washes and stored in 100% methanol at −20°C. FFPE tissue sections were rehydrated through a series of decreasing ethanol washes. Hybridization chain reaction probes were purchased from Molecular Instruments, Inc. Standard protocols for whole-mount mouse embryos and FFPE tissue sections were followed as described by the manufacturer (Molecular Instruments, Inc.). The following probes were used on whole-mount embryos: *Dll1, Sox2, Six2, Meox1, Pax2, Hes7, Tbx18, Osr1, Snai1, Uncx4.1,* and *Wt1*. The following probes were used on FFPE tissue sections: *Pax2, Cdh1, Cdh2, Osr1,* and *Wt1*. Samples were imaged using a Zeiss LSM780 Confocal microscope. When needed, multiple fields of view were taken using the tiling function in Zeiss ZEN Blue software.

### TUNEL assay

FFPE tissue sections were deparaffinized, rehydrated through a series of decreasing ethanol washes, and rinsed in PBS with 0.1% Triton X-100 prior to TUNEL assay. *In Situ* Cell Death Detection Kit, TMR red (Roche 12156792910) manufacturer’s protocol was followed to detect cell death via DNA fragmentation. Images were taken using a Zeiss LSM780 Confocal microscope.

### Proximity Ligation Assay

FFPE tissue sections were deparaffinized and rehydrated through a series of decreasing ethanol washes. Heat induced epitope retrieval was performed using a 20mM sodium citrate pH 6.0 solution in a pressure cooker for 30 minutes followed by pressure release and gradual cooling to 45°C. Sections were washed in PBS with 0.1% Tween-20 prior to following the manufacturer’s protocol. Duolink Proximity Ligation Assay *In Situ* Detection Reagents (Sigma-Aldrich DUO92002, DUO92004, DUO92008) were used according to the manufacturer’s protocol. In brief, sections were blocked followed by overnight incubation with primary antibodies at 4°C. Sections were washed then treated with anti-Mouse MINUS and anti-Rabbit PLUS probes for 1 hour at 37°C. Washes were repeated, followed by incubation with ligase at 37°C for 30 minutes. Slides were washed again then treated with polymerase for signal amplification for 100 minutes at 37°C. Samples were stained with DAPI then washed and coverslips were mounted. Images were taken using a Zeiss LSM780 Confocal microscope. Quantification of puncta was performed in FIJI ImageJ. Confocal images were converted to 8-bit grayscale, thresholding values were calculated for each set of images, followed by use of the Analyze Particles function. One-way ANOVA with Bonferroni correction for multiple comparisons was used to determine statistical significance.

### RT-qPCR

The caudal portion of the embryonic tail was dissected from E9.5 embryos just past the last somite. Samples were lysed and mRNA was extracted according to Qiagen RNeasy kit standard protocol. cDNA was synthesized using 50ng/ul random primers and 50uM Oligo(dT) with SuperScript IV reverse transcriptase (Invitrogen). qPCR was performed using Luna Universal qPCR Master Mix (NEB) with Bio-Rad iQ5 multicolor real-time PCR detection system. Cycling conditions were as follows: step 1, 3 min at 95C; step 2, 10 sec at 95C; step 3; 30 sec at 60C; repeat steps 2 and 3 for 40 cycles. Proper amplification was confirmed using a melting curve. Both *Gapdh* and *Actb* were used as positive controls; *Actb* was found to produce the least amount of variance and was subsequently used for normalization. Two-way ANOVA with Bonferroni correction for multiple comparisons was used to determine statistical significance.

### Coimmunoprecipitation and Western Blot Analysis

NIH3T3 fibroblasts were transfected with either pcDNA3.1-*Fgfr1c-Wt-3xFlag*, pcDNA3.1-*Fgfr1c-FCPG-3xFlag*, or pcDNA3.1-*Fgfr1c-FCSPG-3xFlag* via electroporation and stable lines were selected using the Neomycin resistance cassette co-expressed in the vector. Individual colonies were picked and cultured, and overexpression was verified by RT-qPCR and WB analysis. Cells were maintained in DMEM (Gibco, 11965118) supplemented with 10% HyClone FetalClone III (FCIII) serum (Cytivia, SH30109), 0.5X Penicillin/Streptomycin (Gibco, 15140122), 1X Glutamine (Gibco, 25030081), and 500ug/ml G418 (Gold Biotechnology, G-418). Prior to collection, cells were starved overnight for 18 hours in DMEM containing 0.1% FCIII, then treated with 50ng/ml FGF1 for 5 minutes. Cells were collected and lysed in NP-40/Digitonin lysis buffer containing protease and phosphatase inhibitors (Pierce, A32961) for 30 minutes at 4°C. 300 μg of lysate were incubated with either anti-Flag M2 antibodies conjugated to magnetic beads (Millipore Sigma, M8823) or Pierce Protein A/G magnetic beads (Pierce, 88802) coupled with mouse anti-IgG (CST, 33469) overnight for 18 hours at 4C. Beads were collected and washed six times at 4°C. Remaining supernatant was removed and beads were frozen at −20C and transported on dry ice to the Rockefeller University’s Proteomics Resource Center (RRID:SCR_017797) for label-free quantitative mass spectroscopy. Magnetic beads were dissolved in 40uL 10ng/uL trypsin and proteins were eluted from antibody using partial digestion (4h). Supernatant was removed and subjected to reduction (10mM DTT) and alkylation (30mM IAM) followed by an overnight digestion (trypsin+LysC). Digestions were halted by acidifying (TFA). Peptides were solid-phase extracted and analyzed by LC-MS/MS: 60min gradient (2% to 38%) @900nL/min, 100um/12cm built-in-emitter column, high res./high mass accuracy (Q-Exactive mass spectrometer). The data were processed using ProteomeDiscoverer 1.4 and searched (Mascot) against Uniprot’ss mouse database concatenated with common contaminants which includes FBS related proteins. Data were also analyzed using MaxQuant (Version 2.4.2.0). Proteins quantified at 10-fold greater levels than bait were removed as background. Proteins present in two out of three replicates were counted as positive hits.

### RNA Sequencing Analysis

Seurat v4 was used for single cell data analysis (Hao et al., 2021; Satija et al., 2015). Datasets were normalized via log-fold transformation, followed by variable feature selection and linear scaling based on the top 2000 variable features identified. Linear dimensional reduction was performed using Principal Component Analysis and Elbow Plots were used to identify the number of relevant dimensions. Cells were then clustered using a K-Nearest Neighbor graph and the Louvain algorithm using the first twelve PCA dimensions and a resolution of 1.0. Non-linear dimensional reduction was then performed using Uniform Manifold Approximation and Projection with the first twelve PCA dimensions. Seurat and ggplot2 were used to construct FeaturePlots, DotPlots, and ViolinPlots.

For RNA-seq, mRNA was purified from total RNA using poly-T oligo-attached magnetic beads. After fragmentation, the first strand cDNA was synthesized using random hexamer primers, followed by second strand cDNA synthesis. The library was checked with Qubit and RT-PCR for quantification and bioanalyzer for size distribution detection. Quantified libraries were pooled and sequenced on Illumina platforms. Reference genome and gene model annotation files were downloaded from genome website directly. Index of the reference genome was built using Hisat2 (v2.0.5) and paired-end clean reads were aligned to the reference genome using Hisat2. The mapped reads of each sample were assembled by StringTie (v1.3.3b) in a reference-based approach. FeatureCounts (v1.5.0-p3) was used to count the reads numbers mapped to each gene. Then FPKM of each gene was calculated based on the length of the gene and reads count mapped to this gene. Differential expression analysis was performed using the DESeq2R package (1.20.0). The resulting P-values were adjusted using the Benjamini and Hochberg’s approach for controlling the false discovery rate. Genes with an adjusted P-value <=0.05 found by DESeq2 were assigned as differentially expressed. Gene Ontology (GO) enrichment analysis of differentially expressed genes was implemented by the clusterProfiler R package, in which gene length bias was corrected. GO terms with corrected P-value less than 0.05 were considered significantly enriched by differential expressed genes. We used clusterProfiler R package to test the statistical enrichment of differential expression genes in KEGG pathways. Ggplot2 was used to construct plots.

### Statistical analysis

All statistical analyses were performed and graphs made using Graph Pad (Prism) or RStudio.

### Antibodies

The following antibodies were used for Immunohistochemistry: Rabbit anti-Fgfr1 (1:100, CST 9740), Rabbit anti-Fgfr2 (1:100, CST 23328), Rabbit anti-Myc (1:100, CST 2276), Mouse anti-Myc (1:100, Santa Cruz sc-40), Mouse anti-Flag M2 (1:100, Millipore Sigma F1804), Rabbit anti-DYKDDDDK (1:100, CST 14793), Goat anti-Mouse HRP (1:10,000, Jackson ImmunoResearch 115-035-003), Goat anti-Rabbit HRP (1:10,000, Jackson ImmunoResearch 111-035-003). The following antibodies were used for Immunofluorescence: Rabbit anti-E-Cadherin (1:100, CST 3195), Mouse anti-N-Cadherin (1:100, BDBiosciences 610920), Mouse anti-Ret (1:100, Santa Cruz sc-365943), Rabbit anti-Ki-67 (1:100, CST 12202), Rabbit anti-Myc (1:100, CST 2276), Mouse anti-Myc (1:100, Santa Cruz sc-40), Mouse anti-Flag M2 (1:100, Millipore Sigma F1804), Rabbit anti-DYKDDDDK (1:100, CST 14793), Rabbit anti-EEA1 (1:100, CST 3288), Rabbit anti-Rab7 (1:100, CST 9367), Goat anti-Mouse Alexa Fluor Plus 555, 594, and 647 (1:1,000, ThermoFisher A32727 A32742 A32728), Goat anti-Rabbit Alexa Fluor Pluss 555, 594, and 647 (1:1000, ThermoFisher A32732 A32740 A32733). The following antibodies were used for Proximity Ligation Assay: Rabbit anti-Myc (1:100, CST 2276), Mouse anti-Myc (1:100, Santa Cruz sc-40), Mouse anti-Flag M2 (1:100, Millipore Sigma F1804), Rabbit anti-Src (1:100, CST 2109), Mouse anti-Talin (1:100, Sigma T3287), Rabbit anti-FAK (1:100, Abcam), Mouse anti-Cortactin (1:100, Sigma 05-180), Rabbit anti-αCatenin (1:100, Sigma C2081), Mouse anti-Integrin beta 1 (1:100, Transduction Laboratories 610468), Rabbit anti-Eps15 (1:100, CST 12460), Rabbit anti-Pik3c2a (1:100, CST 12402), Rabbit anti-Pik3r2 (1:100, CST 4292), Mouse IgG1 isotype control (1:100, CST 5415), Rabbit IgG isotype control (1:100, CST 3900). The following antibodies were used for Western blot analysis: Rabbit anti-Fgfr1 (1:1,000, CST 9740), Rabbit anti-Fgfr2 (1:1,000, CST 23328), Rabbit anti-Myc (1:1,000, CST 2276), Mouse anti-Flag M2 (1:1,000, Millipore Sigma F1804), Goat anti-Mouse HRP (1:10,000, Jackson ImmunoResearch 115-035-003), Goat anti-Rabbit HRP (1:10,000, Jackson ImmunoResearch 111-035-003).

